# Cardiovascular symptoms of PASC are associated with trace-level cytokines that affect the function of human pluripotent stem cell derived cardiomyocytes

**DOI:** 10.1101/2024.04.11.587623

**Authors:** Jane E. Sinclair, Courtney Vedelago, Feargal J. Ryan, Meagan Carney, Miriam A. Lynn, Branka Grubor-Bauk, Yuanzhao Cao, Anjali K. Henders, Keng Yih Chew, Deborah Gilroy, Kim Greaves, Larisa Labzin, Laura Ziser, Katharina Ronacher, Leanne M. Wallace, Yiwen Zhang, Kyle Macauslane, Daniel J. Ellis, Sudha Rao, Lucy Burr, Amanda Bain, Benjamin L. Schulz, Junrong Li, David J. Lynn, Nathan Palpant, Alain Wuethrich, Matt Trau, Kirsty R. Short

## Abstract

Globally, over 65 million individuals are estimated to suffer from post-acute sequelae of COVID-19 (PASC). A large number of individuals living with PASC experience cardiovascular symptoms (i.e. chest pain and heart palpitations) (PASC-CVS). The role of chronic inflammation in these symptoms, in particular in individuals with symptoms persisting for >1 year after SARS-CoV-2 infection, remains to be clearly defined. In this cross-sectional study, blood samples were obtained from three different sites in Australia from individuals with i) a resolved SARS-CoV-2 infection (and no persistent symptoms i.e. ‘Recovered’), ii) individuals with prolonged PASC-CVS and iii) SARS-CoV-2 negative individuals. Individuals with PASC-CVS, relative to Recovered individuals, had a blood transcriptomic signature associated with inflammation. This was accompanied by elevated levels of pro-inflammatory cytokines (IL-12, IL-1β, MCP-1 and IL-6) at approximately 18 months post-infection. These cytokines were present in trace amounts, such that they could only be detected with the use of novel nanotechnology. Importantly, these trace-level cytokines had a direct effect on the functionality of pluripotent stem cell derived cardiomyocytes *in vitro*. This effect was not observed in the presence of dexamethasone. Plasma proteomics demonstrated further differences between PASC-CVS and Recovered patients at approximately 18 months post-infection including enrichment of complement and coagulation associated proteins in those with prolonged cardiovascular symptoms. Together, these data provide a new insight into the role of chronic inflammation in PASC-CVS and present nanotechnology as a possible novel diagnostic approach for the condition.

## INTRODUCTION

Globally, at least 65 million individuals are thought to suffer from the post-acute sequelae of COVID-19 (PASC; also known as ‘long COVID’)^1^. The current working definition of PASC by the World Health Organisation defines the condition as “the continuation or development of new symptoms three months after an initial SARS-CoV-2 infection”^2^. Whilst PASC resolves for some individuals within six months of the acute infection, in others symptoms can persist for longer than two years^3^. Consistent with the broad definition of PASC, the condition has been associated with >200 different symptoms across multiple organ systems^1^. At present the cause of PASC remains elusive. However, there is a growing of evidence that PASC is associated with chronic inflammation^4-12^. For example, recently Cervia-Hasler^10^ and colleagues demonstrated that at six months post-infection there was increased complement activation in PASC patients. Increased complement activation was associated with thromboinflammation and potentially drove tissue injury to perpetuate further complement activation^10^. Others have found that individuals with PASC display evidence of ongoing neutrophil activity, B cell memory alterations and building autoreactivity^8^. In PASC patients with unresolved lung injury (as indicated by sustained shortness of breath and abnormal chest radiology) higher monocyte expression of C-X-C motif chemokine receptor 6 (CXCR6) and adhesion molecule P-selectin glycoprotein ligand 1 were observed three to nine months post-acute infection^7^. Finally, several studies have described that eight to ten months post-infection, patients with PASC have continued elevations in proinflammatory cytokine levels^13-15^. Whether these immune perturbations persist in individuals who experience PASC symptoms for a prolonged period of time (i.e. over one year after the acute infection) remains less well defined. However, recent evidence suggests spontaneous release of IFN-γ from peripheral blood mononuclear cells of patients with PASC more than 30 months post-infection^11^. The functional significance of such immunological changes is unclear. Whether these immunological changes may simply be biomarkers of disease, rather than driving PASC symptomology, remains to be determined.

PASC is an incredibly heterogeneous disease and there is a growing recognition that PASC is actually an umbrella term for a variety of different conditions, each with its own unique mechanism of disease and biomarkers^9,16^. For example, Openshaw and colleagues recently demonstrated that while Matrilin-2 (MATN2) and dipeptidyl peptidase like 10 (DPP10) were elevated in individuals with PASC who had gastrointestinal symptoms, complement C1q A chain (C1QA) was elevated in individuals with PASC who had cognitive impairment^5^. Similarly, others have described at least two different subtypes of PASC; only one of which is associated with chronic inflammation^8^. Recent data suggest that circulating fibroblast growth factor 21 (FGF21) is elevated in the blood of PASC patients with cognitive deficits but not in those reporting physical deficits^17^. Together, these data suggest that further mechanistic insights into PASC can be obtained by separately considering the different pathotypes, rather than treating PASC as a blanket term for a single unified condition.

One of the most common manifestations of PASC is cardiovascular disease^18^. The American College of Cardiology Solution Set Oversight Committee has defined two separate subsets of cardiovascular disease in PASC^19^. PASC cardiovascular syndrome (PASC-CVS) is a heterogeneous disorder that may include a range of self-reported cardiovascular symptoms (e.g. chest pain, heart palpitations and post-exertional malaise)^19^. In contrast, PASC cardiovascular disease (PASC-CVD) includes more objective signs of cardiovascular disease such as myocarditis, ischemia and arrhythmia^19^. PASC-CVS is particularly difficult to consistently diagnose, given that diagnosis is dependent on self-reported symptoms. Accordingly, the incidence of PASC-CVS varies between studies. However, at 5 months post-hospital discharge for COVID-19 between 21-28% of individuals reported chest pain and palpitations^20^. It has been hypothesised that ongoing inflammation may drive these cardiovascular symptoms^21,22^. For example, a proinflammatory response may result in the destabilization of the atherosclerotic plaques leading to embolisms, ischemia and tissue damage^22^. However, direct evidence for persistent inflammation in individuals with the cardiovascular complications of PASC is sparse. Individuals with severe post-exertional malaise approximately 400 days post-SARS-CoV-2 infection had a proteomic signature consistent with a persistent inflammatory state and endothelial dysfunction^23^. Similarly, individuals with persistent cardiorespiratory symptoms (defined as dyspnoea, heart palpitations and/or chest pain) had elevated levels of interleukin (IL) 1 receptor 2 approximately 6 months after hospitalisation for SARS-CoV-2^5^. There is thus a clear need to better understand the relationship between inflammation and PASC-CVS, in particular at prolonged periods (i.e. >1 year) post SARS-CoV-2 infection.

Here, we obtained blood samples >1-year post-infection from individuals with PASC-CVS (as defined by self-reported symptoms of chest pain and/or heart palpitations persisting for >3 months after SARS-CoV-2 infection). We showed that compared to samples obtained from individuals with a resolved SARS-CoV-2 infection (and no persistent symptoms i.e. ‘Recovered’), individuals with PASC-CVS had a blood transcriptomic signature associated with inflammation. This was coupled with elevated levels of pro-inflammatory cytokines. However, these cytokines were present in trace amounts, such that they could only be detected with the use of novel nanotechnology^24^. Despite the differences in cytokines being trace level, these differences were sufficient to have a direct effect on the functionality of pluripotent stem cell derived cardiomyocytes *in vitro*. Proteomics demonstrated further differences in plasma from PASC-CVS and Recovered patients including enrichment of complement and coagulation associated proteins. Together, these data provide a new insight into the long-term cardiovascular complications of COVID-19.

## MATERIALS AND METHODS

### Ethics & Participants

Participants with PASC were defined as individuals with persistent symptoms >12 weeks post-infection who reported ongoing discomfort at the time of sample collection.

Participants with PASC-CVS were defined as individuals who experienced chest pain and/or chest tightness after SARS-CoV-2 infection >12 weeks post-infection ‘Recovered individuals’ were defined as those who were infected with SARS-CoV-2 but did not report persistent symptoms >12 weeks post-infection.

‘Healthy individuals’ were defined as individuals who were recruited prior to community transmission of SARS-CoV-2 in Queensland or South Australia (i.e. prior to the end of 2021).

Details of the individual cohorts are provided below.

#### PASC cohort 1, PASC-CVS cohort 1 and Recovered cohort 1

Participants for the South Australia cohort of the study were enlisted through the Central Adelaide Health Network (CALHN), with the research protocol receiving approval from the CALHN Human Research Ethics Committee in Adelaide, Australia (Approval No. 13050). Inclusion criteria involved a PCR-confirmed SARS-CoV-2 infection from nasopharyngeal swabs, occurring in March and April of 2020 for all participants, the capability to attend follow-up visits and the provision of voluntary informed consent. Analyses of blood samples collected from this cohort at 12, 16 and 24 weeks post-infection have been previously reported^12^. For the current study, blood samples were additionally obtained at approximately 44- and 68-weeks post-infection. Convalescent patients were asked to complete a retrospective questionnaire detailing self-reported symptoms related to PASC (listed as Additional File 10 in ^12^) at approximately 18 months (mean 70.4 weeks, min 61 weeks, max 74 weeks) post-infection. Furthermore, clinical data were obtained to identify convalescent individuals referred to a long COVID clinic operated by the South Australian State health service (SA Health). Blood (54 ml/individual) was collected in serum separator tubes (acid citrate dextrose (ACD)) or ethylenediaminetetraacetic acid (EDTA) tubes and processed for plasma isolation. A 2.5 ml blood sample for RNA sequencing was collected into PAXgene® tubes (762165 BD, North Ryde, Australia) and stored at -80°C until processing.

#### PASC-CVS cohort 2 & Recovered cohort 2

Individuals who reported a laboratory-confirmed case of SARS-CoV-2 infection in Queensland (as defined by nasopharyngeal swab or serology) were recruited from the COVID OZGenetics study (University of Queensland Human Research Ethics approval UQ 2020001490). Informed consent was obtained from all subjects. All recruited participants reported a SARS-CoV-2 infection between 4/2/2020 and 4/9/2020. Participants completed online self-report questionnaires developed in-house. Questionnaires included self-reported information on the date of testing positive, pre-COVID-19 activities, COVID-19 experiences and COVID-19 post-experience. A lifestyle and environmental exposure questionnaire used across multiple studies operated by the University of Queensland’s Human Studies Unit (HSU) which included a full self-report medical history was also administered to all participants. Participants were asked to report on seven different symptoms across multiple time frames (Brain Fog; Chest tightness; Fast Heart rate; Fatigue; Loss of ability to smell/taste; Shortness of breath). Blood samples were collected in VACUETTE® (Griener Bio-One GmbH, Austria) EDTA and SST tubes. Plasma and serum were isolated from respective tubes by centrifugation at 3000rpm for 10mins with low brake with transit times not exceeding 72 hours from time of collection. Plasma fractions of 1.3mLs were stored at - 80°C. Where relevant, an aliquot of whole blood was taken from an EDTA tube and added to RNALater.

#### PASC-CVS cohort 3

Ethical approval to collect blood samples from individuals with PASC was granted by the Mater Misericordiae Ltd Human Research Ethics Committee (MML HREC) under project reference HREC/MML/89069 (V2) and accepted by QIMR Berghofer Human Research Ethics Committee under project reference p3844. All participants provided written informed consent. Participants were administered a PASC survey which included date of SARS-CoV-2 infection and a description of symptoms which persisted >3 months post-infection (including fatigue, shortness of breath, brain fog, heart palpitations, chest pain and loss of taste/smell) across different time frames. Blood was either collected in EDTA or heparin tubes (BD Biosciences, NJ, USA) or BD SST™ Vacutainers® (BD Biosciences). To isolate plasma, tubes containing blood samples were spun down at 1500[*g* for 10 min and stored at -80°C until processing.

#### Healthy Cohort 1

Unexposed healthy participants were recruited from South Australia (CALHN Human Research Ethics Committee in Adelaide, Australia (Approval No. 13050). Healthy controls had no respiratory disease, no positive COVID-19 PCR test in 2020/21, no known significant systemic diseases, and negative anti-Spike and anti-RBD serology

#### Healthy Cohort 2

Unexposed healthy participants were recruited from the Red Cross in Queensland (University of Queensland Human Research Ethics approval UQ 2020001490 & Bellberry Human Ethics Research Committee 2015-12-817-A-6). Alternatively, healthy volunteers were recruited from the Mater Research Institute (Mater Misericordiae Ltd Human Research Ethics Committee 17/MHS/78. Informed consent was obtained from all subjects. Blood was either collected in EDTA or heparin tubes (BD Biosciences, NJ, USA) or BD SST™ Vacutainers® (BD Biosciences). To isolate plasma, tubes containing blood samples were spun down at 2000[*g* for 10 min and stored at -80°C until processing.

### Human cardiomyocyte differentiation

All human iPSC studies were carried out with consent from The University of Queensland’s Institutional Human Research Ethics approval (HREC#: 2015001434). WTC-11 hiPSC cells (Gladstone Institute of Cardiovascular Disease, UCSF)^25,26^ were maintained in mTeSR Plus medium with supplement (Stem Cell Technologies, Cat.#05825) and cultured on Vitronectin XF (Stem Cell Technologies, Cat.#07180) coated plates (Nunc, Cat.#150318) at 37°C with 5% CO_2_. HiPSC-derived cardiomyocytes were generated according to a modified monolayer platform protocol as previously described^27,28^. Briefly, on day -1 of differentiation, hiPSC were dissociated using 0.5% EDTA, plated onto Vitronectin XF-coated flasks (Nunc, Cat.#156367) at a density of 1.12 × 10^5^ cells/cm^2^, and cultured overnight in mTeSR Plus medium supplemented with 10 µM Y-27362 ROCK inhibitor (Stem Cell Technologies, Cat.#72308). Differentiation was induced on day 0 by first washing cells with phosphate-buffered saline (PBS) when the monolayer reached approximately 80% confluence, then changing the culture medium to RPMI 1640 Medium (ThermoFisher, Cat.#11875119) containing 3 μM CHIR99021 (Stem Cell Technologies, Cat.#72054), 500 μg/mL bovine serum albumin (BSA, Sigma Aldrich, Cat.#A9418), and 213 μg/mL ascorbic acid (AA, Sigma Aldrich, Cat.#A8960). After 3 days of culture, the medium was replaced with RPMI 1640 containing 5 μM XAV-939 (Stem Cell Technologies, Cat.#72674), 500 μg/mL BSA, and 213 μg/mL AA. On day 5, the medium was exchanged to RPMI 1640 containing 500 μg/mL BSA, and 213 μg/mL AA without additional supplements. From day 7 and onward, the cultures were fed every other day with RPMI 1640 containing 2% B27 supplement with insulin (Life Technologies Australia, Cat.#17504001). Starting on days 9 and 11 of differentiation, spontaneous beating cardiomyocytes were typically observed.

### RNA extraction

RNA extraction and genomic DNA elimination from whole blood was carried out using the PAXgene® Blood RNA kit (762164, Qiagen, Feldbachstrasse, Germany) as per the manufacturer’s instructions (Cohort 1). Alternatively, RNA was extracted using the Ribopure Blood RNA Kit (Invitrogen™) (Cohort 2). Cardiomyocyte RNA was extracted using the Qiagen RNeasy Plus Kit according to the manufacturer’s instructions.

### RNA Seq

#### Whole Blood – PASC, PASC-CVS, Recovered and Healthy cohort 1

Total RNA was converted to strand-specific Illumina compatible sequencing libraries using the Nugen Universal Plus Total RNA-Seq library kit from Tecan (Mannedorf, Switzerland) as per the manufacturer’s instructions (MO1523 v2) using 12[cycles of PCR amplification for the final libraries. An Anydeplete probe mix targeting both human ribosomal and adult globin transcripts (HBA1, HBA2, HBB, HBD) was used to deplete these transcripts. Sequencing of the library pool (2 × 150 bp paired-end reads) was performed using 2 lanes of an S4 flowcell on an Illumina Novaseq 6000. Sequence read quality was assessed using FastQC version 0.11.4 and summarised with MultiQC version 1.8 prior to quality control with Trimmomatic version 0.38. Reads that passed all quality control steps were then aligned to the Human genome (GRCh38 assembly) using HISAT2 version 2.1.0. The gene count matrix was generated with FeatureCounts version 1.5.0-p2 using the union model with Ensembl version 101 annotation. The count matrix was then imported into R version 4 0.3 for further analysis and visualisation in ggplot2 v2.3.3. Counts were normalised using the trimmed mean of M values (TMM) method in EdgeR version 3.32 and represented as counts per million (cpm) Svaseq v3.38 was applied to remove batch effects and other unwanted sources of variation in the data. Differential gene expression analysis was performed using the glmLRT function in EdgeR adjusting for sex and batch (run) in the model. Genes with <[3[cpm in at least 15 samples were excluded from the differential expression analysis. Pathway and Gene Ontology (GO) overrepresentation analysis was carried out in R using a hypergeometric test. Gene Set Enrichment Analysis (GSEA) was carried out using the camera function in the EdgeR library with the Molecular Signatures Database (MSigDB) R package (msigdbr v7.4.1). Blood transcriptional module (BTM) analysis was carried using a pre-defined set of modules defined by Li et al. as an alternative to pathway-based analyses^29^.

#### Whole blood – PASC-CVS and Recovered cohort 2

Library preparation and sequencing was performed at the University of Queensland Sequencing Facility. RNA-Seq libraries were prepared using the Illumina stranded total RNA prep ligation with Ribo-Zero plus kit (Illumina, 20040529) and IDT for Illumina RNA UD Indexes (illumina, 20040554) according to the standard manufacturer’s protocol (illumina, Document # 1000000124514 v03, June 2022) The libraries were quantified on the Perkin Elmer LabChip GX Touch with the DNA High Sensitivity Reagent kit (Perkin Elmer, CLS760672). Libraries were pooled in equimolar ratios. Sequencing was performed using the Illumina NovaSeq 6000. The library pool was diluted and denatured according to the standard NovaSeq protocol and sequenced to generate paired-end 152 bp reads using a NovaSeq 6000 SP reagent kit v1.5 (300 cycles) (Illumina, 20028400). After sequencing, fastq files were generated using bcl2fastq2 (v2.20.0.422), which included trimming the first cycle of each insert read due to expected low diversity. RNA-Seq raw read quality was evaluated with FastQC (v0.12.1) ^30^ prior to quality control with Trimmomatic v0.39^31^. Reads that passed all quality control steps were then aligned to the GRCh38 human genome with HISAT2 (v2.2.1)^32^. A gene count matrix was generated with FeatureCounts (v2.0.6, union model) ^33^ and the Ensembl v109 annotation. The count matrix was then imported into R (v4.2) for further analysis. Counts were normalised using the trimmed mean of M values (TMM) method in EdgeR (v3.38)^34^ and represented as counts per million (cpm). Gene set enrichment analysis was carried out with the fgsea R package (v1.22)^35^ as described above.

#### Cardiomyocytes

Library preparation and sequencing was performed at the University of Queensland Sequencing Facility. RNA-Seq libraries were generated using the Illumina Stranded mRNA Library Prep Ligation kit (illumina, 20040534) and IDT for Illumina RNA UD Indexes (illumina, 20040553) according to the standard manufacturer’s protocol. Libraries were pooled in equimolar ratios. Sequencing was performed using the Illumina NovaSeq 6000. The library pool was diluted and denatured according to the standard NovaSeq protocol and sequenced to generate paired-end 102 bp reads using a 100 cycle NovaSeq reagent Kit v1.5 (illumina, 20028401). After sequencing, fastq files were generated using bcl2fastq2 (v2.20.0.422), which included trimming the first cycle of each insert read due to expected low diversity. RNA-Seq raw read quality was evaluated with FastQC (v0.12.1) ^30^ prior to quality control with Trimmomatic v0.39^31^. Reads that passed all quality control steps were then aligned to the GRCh38 human genome with HISAT2 (v2.2.1) ^32^. A gene count matrix was generated with FeatureCounts (v2.0.6, union model) ^33^ and the Ensembl v109 annotation. The count matrix was then imported into MATLAB for further analysis. The histogram of raw log(counts) was used to determine the filter threshold for count of gene expression. Any gene expression falling below the filter threshold was taken as zero. Batch effect removal was performed using ComBat factor analysis in MATLAB with the model matrix defined using ‘batch’ and ‘class’ as regressor variables. Bioconductor differential expression analysis was performed using the rnadeseq.m function in MATLAB on the batch-corrected gene expression counts. Gene set enrichment analysis was carried out with the fgsea R package (v1.22) ^35^ as described above.

### LEGENDplex assay

The LEGENDplex Human Anti-Virus Response Panel (IL-1β, IL-6, IL-8, IL-10, IL-12p70, IFN-α2, IFN-β, IFN-λ1, IFN-λ2/3, IFN-γ, TNF-α, IP-10 and GM-CSF; BioLegend) assay was performed on patient plasma samples as per the manufacturer’s instructions.

### ELISA

The Human Monocyte Chemotactic Protein 1 (MCP-1) ELISA Kit (MyBioSource) assay was performed on patient plasma samples as per the manufacturer’s instructions.

### Immunostorm chip and cardiac chip

The chips were prepared and used as previously described^24^. For the immunostorm chip, the nanopillar surface of the chip was functionalised with a cocktail of antibodies against IL-1β (AF-201, polyclonal), IL-6 (MAB9540, clone 973132), IL-12p70 (MAB611R, clone 24945R), and MCP-1 (AF-279, polyclonal) obtained from R&D Systems. Surface enhanced Raman scattering (SERS) nanotags were conjugated with antibodies against IL-1β (MAB601, clone 2805), IL-6 (AF-206, polyclonal), IL-12p70 (MAB611, clone 24945), IL-12p70 (MAB219, clone 24910), or MCP-1 (MAB679, clone 23007) obtained from R&D Systems. For patient sample analysis using the immunostorm chip 10 µL of diluted human plasma (1:10 dilution with 1x PBS) was used.

For the cardiac chip, the nanopillar surface was conjugated with antibodies against B-Type Natriuretic Peptide (BNP) (ab236101, polyclonal, Abcam), Atrial natriuretic peptide (ANP) (MA5-37730, polyclonal, Invitrogen), MIP-1β (PA5-34509, polyclonal, Invitrogen), cardiac troponin I (cTnI) (MA1-22700, polyclonal, Invitrogen) and Creatine Kinase MB (CKMB) (PA5-28920, polyclonal, Invitrogen). SERS nanotags were conjugated with antibodies against BNP (ab236101, polyclonal, Abcam), ANP (MA5-37730, polyclonal, Invitrogen), MIP-1β (PA5-34509, polyclonal, Invitrogen), cTnI (MA1-20112, clone 16A11, Invitrogen) and CKMB (PA5-28920, polyclonal, Invitrogen). For patient sample analysis using the cardiac chip 10 µL of diluted human plasma (1:50 dilution with 1x PBS) was used.

### Proteomics

#### Proteomic Sample Preparation

2 µL of nondepleted whole plasma was added to 98 µL of 50 mM Tris-HCl buffer pH 8, 6 M guanidine hydrochloride, and 20 mM dithiothreitol (DTT) and incubated for 30 min at room temperature. Cysteines were alkylated by the addition of acrylamide to a final concentration of 50 mM and incubation at 30 °C for 1 h with shaking. Excess acrylamide was quenched with the addition of DTT to a final additional concentration of 10 mM. Samples were transferred to 10 kDa amicon cut-off filter columns and were washed three times with the addition of 500 µL 50 mM NH_4_HCO_3_ and centrifugation at 14,000 g for 20 min. Following the washes, an additional 150 µL NH_4_HCO_3_ and 2 µg of sequencing grade porcine trypsin were added, and samples were digested at 37 °C for 16 h in an enclosed humidified incubator. Digested peptides were eluted by centrifugation at 10,000 *g* for 10 min and a final wash of 50 µL 50 mM NH_4_HCO_3._ Peptides were desalted and concentrated with C18 ZipTips (Millipore), dried down with vacuum centrifugation and resuspended in 20 µL 0.1 % formic acid (FA).

#### Mass Spectrometry

Proteomic samples were analysed in randomised injection order by data-independent acquisition (DIA) using a ZenoTOF 7600 mass spectrometer (SCIEX) coupled to a Waters M-Class UPLC system. 400 ng of sample was injected onto a Waters nanoEase M/Z HSS T3 C18 column (300 µm, 150 mm, 1.8 µm, 100 Å). Peptides were separated with buffer A (0.1 % FA in water) and buffer B (0.1% FA in acetonitrile). Chromatography was performed at 5 µL/min with the column at 40 °C, with LC program: 0 – 0.5 min, 1 % B; 0.5 – 0.6 min, linear gradient to 5 % B; 0.6 – 22 min, linear gradient to 35 % B; 22 – 23 min, linear gradient to 60 % B; 23 – 23.4 min, linear gradient to 90% B; held at 90% B for 3 min; re-equilibration in 1% B for 1.6 min. Eluted peptides were directly analysed on a ZenoTOF 7600 instrument (SCIEX) using an OptiFlow Micro/MicroCal source. Curtain gas, 35 psi; CAD gas, 7; Gas 1, 20 psi; Gas 2, 15 psi; source temp, 150 °C; spray voltage, 5000 V; DP, 80 V; CE, 10 V. MS1 spectra were acquired at *m/z* 400-1500 (0.1 s) followed by DIA with 65 variable windows across an *m/z* range of 400-850 with fragment data acquired across 140 – 1750 *m/z* (0.013 s) with Zeno pulsing on and threshold set to 100,000 cps. Variable window sizes were generated using the SWATH Variable Window Calculator (SCIEX), based on a representative pooled plasma sample. Parameters included 1 Da window overlap, and a minimum window width of 5 Da. Dynamic collision energy was automatically assigned by the Analyst software (SCIEX) based on variable window *m/z* ranges.

#### Data Analysis

A spectral library was generated by processing all raw DIA files with DIA-NN (v1.8)^36^. A library-free search was performed using a FASTA file containing 20,428 human proteins (Uniprot reviewed, downloaded 7 December 2023) and common mass spectrometry contaminants. Default settings for a library-free search were used with the following changes: propionamide was set as a fixed modification on cysteine residues; N-term M excision was not included; minimum peptide length, 4; precursor charge range, 2-5; protein inference, Genes; quantification strategy, Robust LC. The spectral library generated by DIA-NN was converted into a PeakView (SCIEX) compatible format using a custom-designed script. The PeakView (v2.2, SCIEX) SWATH MicroApp was used to determine the abundance of peptide fragments, peptides and proteins as previously described^37^. Protein abundances were recalculated using a 1% false discovery rate cut-off. Normalisation was performed to total human protein abundance in each sample.

### Treatment of hiPSC cardiomyocytes

Differentiated cardiomyocytes were replated on day 15 of differentiation for contractility recording using the CardioExcyte 96 system (Nanion Technologies GmbH). Cells were seeded at a density of 50,000 cells/well onto Vitronectin XF-coated NSP-96 plates (Nanion Technologies, Munich) and transferred to the CardioExcyte 96 platform for baseline monitoring. Upon stabilisation of baseline readings for human plasma experiments plasma was diluted 1:40 in media (RPMI [Thermo Fisher Scientific, U.S.A.] with B27 Supplement [Thermo Fisher Scientific, U.S.A.]), then transferred to the hiPSC-derived cardiomyocytes for 48 hours of treatment either in the presence or absence of 100ng/mL of dexamethasone (Sigma-Aldrich, U.S.A). For cytokine spiking experiments, recombinant cytokines IL-12, IL-1B, IL-6 and MCP-1 (Abcam, Australia) were diluted in media at the stipulated concentrations. Contractility recordings performed with CardioExcyte 96 were obtained with 1 ms time resolution and 1 kH sampling rate. All experiments were conducted under physiological conditions using a built-in incubation chamber in the system (37 °C, 5% CO2 and 80% humidity). The CardioExcyte NSP-96 plates utilise two gold electrodes on a rigid surface in each well to study physiological changes in contractility via continuous impedance measurements. The readouts for detection are amplitude, beat rate, upstroke velocity, and relaxation velocity. For recombinant cytokine experiments data were displayed a raw data, while experiments using patient plasma display data as relative to a standard media control treatment.

### Statistics and data availability

Data were analysed on Graphpad v9.0.2 (Dotmatics). Normal distribution of data was assessed with the Shapiro-Wilk test. Statistical significance was determined with a Kruskal-Wallis test with Dunn’s multiple comparison test, Mann-Whitney U test or Welch ANOVA test and Dunnett’s multiple comparison test where p<0.05 was considered significant. Alternatively, ANCOVAs were performed using MATLAB and the associated code is available at https://doi.org/10.48610/b3182cf. Adjustments to p-values for multiple testing were made using the Benjamini-Hochberg procedure. The processed mass spectrometry proteomics data are available at ProteomeXchange Consortium via the PRIDE partner repository with the dataset identifier PXD050202. RNASeq data is available upon request from the Gene Expression Omnibus.

## RESULTS

To investigate the long-term effects of SARS-CoV2 infection on the blood transcriptome, individuals were recruited from Cohort 1 (South Australia) at approx. 44 weeks (∼308 days) and 68 weeks (∼476 days) post-infection (wpi, Supplementary Table 1) and total bulk RNA sequencing was performed on whole blood (mean 53.2 million 2x150bp read pair per sample sequenced). This cohort consisted of individuals who reported persistent symptoms more than 12 wpi (PASC, n=25), those whose symptoms did not persist long-term (Recovered, n=11). RNA sequencing was also performed on samples from contemporaneous healthy controls (n = 14) with negative serology for the SARS-CoV-2 Spike protein (Supplementary Table 1). We previously reported on alterations in the transcriptome of this cohort at 12, 16 and 24 wpi ^12^. UMAP analysis of the gene expression data revealed that samples collected from healthy controls (HC) separated from those with any past COVID-19 (‘Any COVID-19’). However, samples from PASC/Recovered individuals tended to cluster together (Figure 1A). Interestingly, there were approximately twice as many DEGs identified in PASC (>400 DEGs) compared to Recovered individuals (<200 DEGs) when both groups were compared to HCs at 44wpi (Figure 1B). A similar number of DEGs were identified in PASC and Recovered individuals at 68 wpi (∼200 DEGs) relative to Healthy Controls (Figure 1B). To investigate which pathways, modules or biological processes were enriched among the DEGs, we performed a comprehensive gene set enrichment analysis (GSEA) using the molecular signatures database (mSigDB), focusing on gene-sets from the KEGG, REACTOME and Gene Ontology (GO) databases together with pre-defined blood transcriptional modules (BTMs)^29^. Compared to HCs, any COVID-19 individuals showed an up-regulation of pathways related to RNA processing, transcription and translation, signatures which were driven by the upregulation of ribosomal RNA (rRNA genes). (Figure 1C). We then applied GSEA to compare PASC and Recovered individuals. Relative to Recovered individuals, PASC individuals had an up-regulation of the same translation and ribosomal signatures observed in any COVID-19 individuals at 44 wpi, but not 68 wpi (Figure 1D). At 68 wpi, the most significantly enriched Reactome pathway and Gene Ontology terms were related to erythrocytes and the production of antimicrobial peptides (Figure 1D). T cell and NK cell related BTMs were also found to be up-regulated in PASC compared to Recovered individuals (Figure 1D).

**Figure 1:**
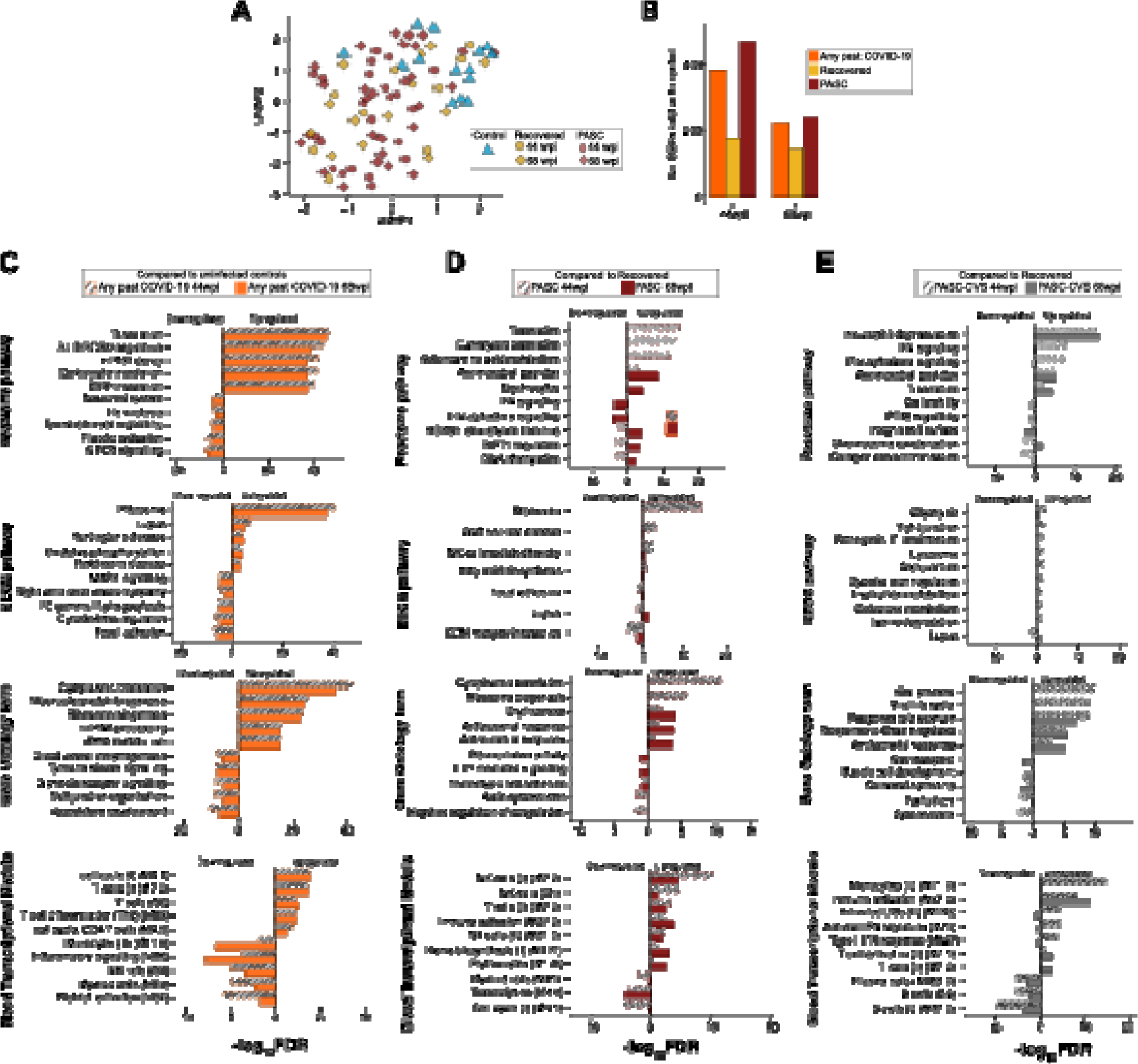
Transcriptomic analyses of PASC, PASC-CVS, Recovered and Healthy blood samples from Cohort 1. **(A)** UMAP of whole blood gene expression data at 44 and 68-weeks post infection (wpi) for PASC (n=25) and recovered (n=11) participants and healthy controls (n=14). **(B)** The number of differentially expressed (DE) genes (FDR <1_0.05 and fold change >1_1.5-fold) identified at each timepoint. **(C)** Topmost up- and downregulated pathways and modules ranked by False Discovery Rate (FDR) in those with past COVID-19 relative to uninfected controls. (**D**) Topmost up- and downregulated pathways and modules ranked by FDR in PASC relative to recovered. (**E)** Topmost up- and downregulated pathways and modules ranked by FDR in PASC-CVS relative to recovered.

A subset of the PASC group was considered as PASC-CVS based on self-reported chest pain (n=4 at each timepoint) (Figure 1E). Compared to Recovered individuals, PASC-CVS individuals had a more pronounced up-regulation inflammation related pathways/terms including the neutrophil degranulation, antimicrobial peptides, and response to bacterium pathways/terms at both 44 and 68 wpi and interferon signalling and at 44 wpi only (Figure 1E). BTMs related to monocytes, DCs, and immune activation were also up-regulated (Figure 1E). Many of these pathways were also significant when comparing PASC-CVS to other PASC individuals (Figure 1D&E).

To determine whether the observed transcriptional differences in the blood of PASC-CVS donors could be replicated in an independent cohort, blood samples were additionally collected from age and sex-matched PASC-CVS (n=5) and Recovered (n=4) participants from Cohort 2 at >520 days post infection (Supplementary Table 2). Bulk RNASeq was independently performed on these whole blood samples (mean 55.9 million 2x150bp reads per sample). GSEA with pre-defined BTM identified numerous pathways were significantly enriched in PASC-CVS samples, predominately those associated with the immune response (e.g. ‘neutrophil degranulation’, ‘humoral immune response’ ‘interferon alpha/beta signalling’ and ‘inflammatory molecules’) (Figure 2). Pathways such as ‘platelet activation’ and blood coagulation were also enriched in PASC-CVS samples (Figure 2).

**Figure 2:**
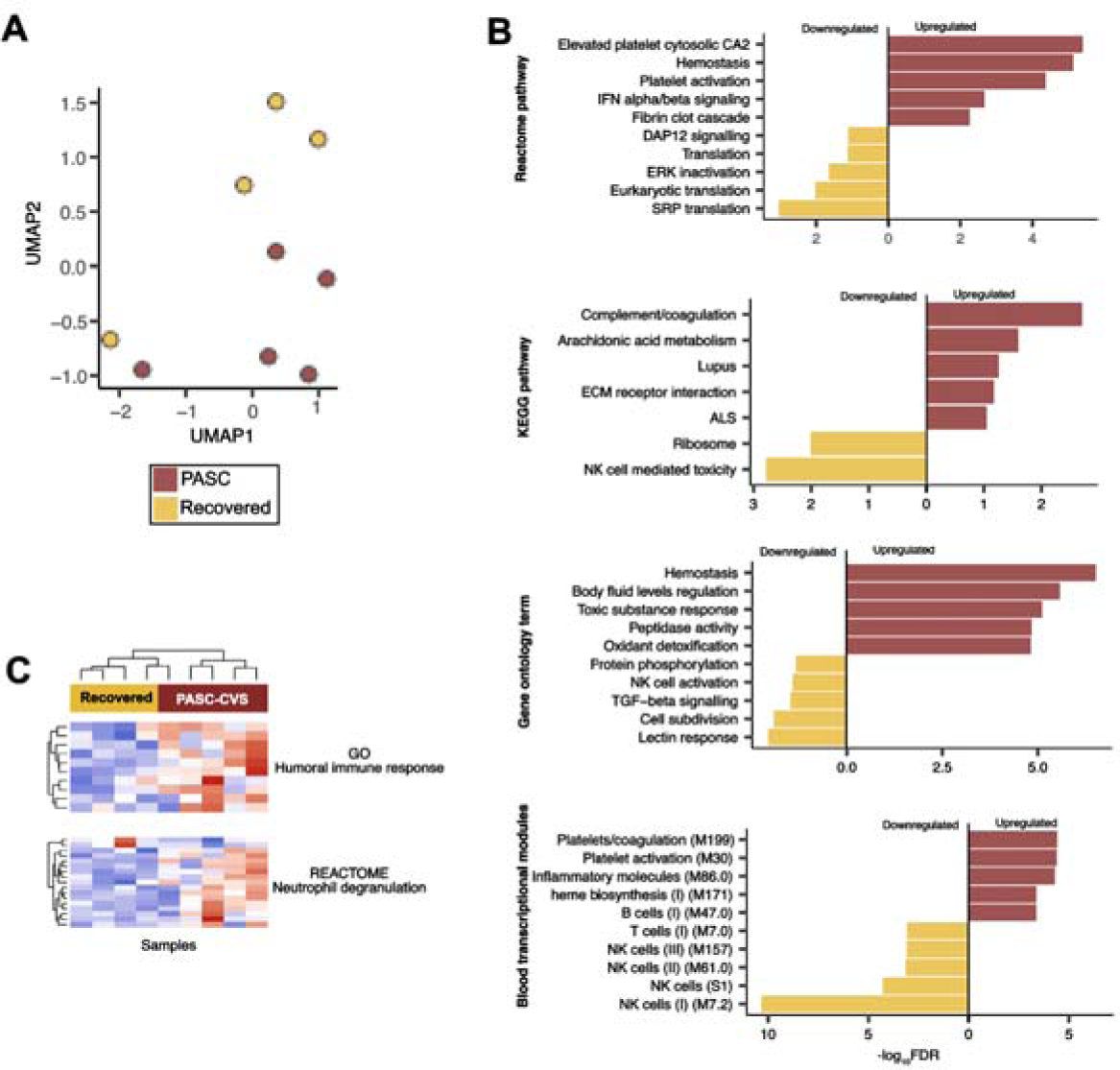
Transcriptomic analyses of PASC-CVS and Recovered blood samples in Cohort 2. **(A)** UMAP of whole blood gene expression for PASC-CVS (n=5) and recovered (n=4) participants. **(B)** Topmost up- and downregulated pathways and modules ranked by FDR in PASC-CVS relative to recovered. **(C)** Heatmap showing the expression of genes increase among PASC-CVS in the GO Humoral immune response (GO:0006959), and REACTOME neutrophil degranulation pathway (R-HSA-6798695) pathways.

In light of differences in inflammation associated transcripts in the blood PASC-CVS donors from two independent cohorts, we created a larger PASC-CVS cohort that included participants from South Australia (Cohort 1), the University of Queensland (Cohort 2) and the Mater Hospital (Cohort 3) (Table 1; Supplementary Figure 1). Recovered individuals were contemporaneously recruited from South Australia (Cohort 1) and the University of Queensland (Cohort 2) (Table 1; Supplementary Figure 1). Individuals with PASC-CVS were defined as those with chest pain and/or heart palpitations. Whilst some individuals had these symptoms exclusively, others reported several additional symptoms including shortness of breath, brain fog and fatigue (Supplementary Figure 1). Recovered individuals provided blood samples approximately 476 days post-SARS-CoV-2 infection (minimum time 289 days; maximum time 599 days). Similarly, individuals with PASC-CVS provided blood samples approximately 497 days post-SARS-CoV-2 infection (minimum time 224 days; maximum time 623 days). As previously described^38^ individuals with PASC-CVS had a lower incidence of mild acute disease but a higher incidence of moderate acute disease compared to Recovered individuals (Table 1). There was no significant difference in the incidence of severe acute disease between the two patient groups (Table 1). Importantly, there were significant differences between groups in terms of age, sex distribution and site (Table 1). Accordingly, subsequent analyses between Recovered and PASC-CVS donors adjusted for these covariates.

**Table 1:** Participant characteristics.

|  | **Recovered** | **PASC-CVS** | **P-value** |
| --- | --- | --- | --- |
| Total | 41 | 23 |  |
| Mean days post-infection that a blood sample was acquired (mean ± SD) | 465 ± 78 | 497 ± 86 | 0.459<sup>1</sup> |
| Sex (M;F) | 22;19 | 5;18 | 0.018<sup>2</sup> |
| Age (mean ± SD) | 56.21±13.8 | 48.37 ±11.30 | 0.024<sup>3</sup> |
| *Severity of acute infection*<sup>4</sup> |  |  | 0.005<sup>2</sup> |
| Mild | 34 | 10 |  |
| Moderate | 6 | 11 |  |
| Severe | 1 | 1 |  |
| *Recruitment site* |  |  | 0.05<sup>2</sup> |
| Cohort 1 (South Australia) | 20 | 6 |  |
| Cohort 2 (University of Queensland) | 21 | 15 |  |
| Cohort 3 (Mater) | 0 | 3 |  |

<sup>1</sup>Mann-Whitney Test <sup>2</sup>Chi square test <sup>3</sup>Student's t-test <sup>4</sup>Data from one PASC-CVS participant was not available

Considering the transcriptional differences between PASC-CVS and Recovered donors included pathways related ‘immune activation’ and ‘cytokine mediated signalling pathway’ (Figures 1 & 2) we assessed levels of chemokines, cytokines and interferons in participant plasma using a multiplex bead-based assay (Supplementary Figure 2) or ELISA (MCP-1; Supplementary Figure 2). There was no significant difference in the means of any of the assessed cytokines/chemokines between the Recovered and PASC-CVS samples, except for a statistically significant difference in the levels of IFN-λ1 (Supplementary Figure 2; Recovered adjusted mean = 21.1pg/mL & PASC-CVS adjusted mean = 10.12pg/mL) and IFN-α2 (Supplementary Figure 2; Recovered adjusted mean = 2.58pg/mL & PASC-CVS adjusted mean = 1.48pg/mL). Importantly, many samples were at or below the detection limit of the assay (Supplementary Figure 2). As few cytokines were detected in donor plasma using conventional technology, we sought to employ a novel nanotechnology - the immunostorm chip^24^ - to detect trace-level cytokines. The immunostorm chip combines an array with designated gold-topped nanopillars and plasmonic barcodes for SERS to achieve single molecule sensitivity (Supplementary Figure 3). Specifically, the immunostorm chip was designed to detect trace-levels of IL-1β, IL-6, IL-12 and MCP-1 (Supplementary Figure 3). Strikingly, there were significantly increased levels of IL-12, IL1β, MCP-1 and IL-6 in plasma of PASC-CVS donors compared to Recovered donors (Figure 3) (adjusted mean difference: 309.45fg/mL [IL-12]; 72.86 fg/mL [IL-1β]; 48.3 fg/mL [MCP-1] & 127.96 fg/mL [IL-6]) (Figure 3). In PASC-CVS donors there were strong positive correlations between the levels of IL-12, IL-12 and IL-1β (Supplementary Figure 4). MCP-1 levels were weakly correlated with the other cytokines measured and there was a negative correlation between MCP-1 levels and time since infection (Supplementary Figure 4).

**Figure 3:**
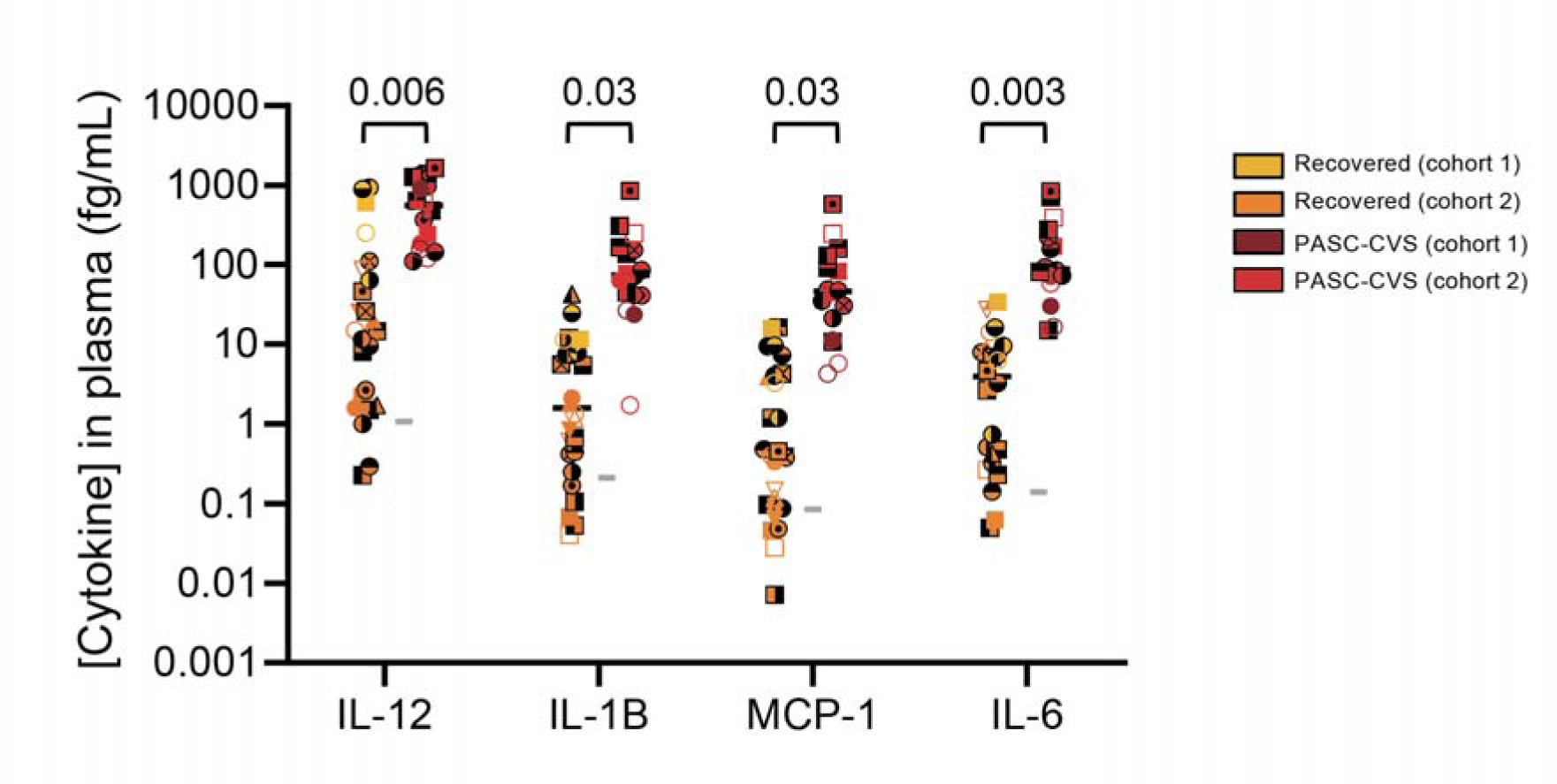
Nanotechnology can detect elevated cytokine levels in the plasma of individuals with PASC-CVS. Cytokine levels detected in the plasma of participants by immunostormchip. Median is shown in all graphs by a black bar. Statistical significance was determined with an ANCOVA adjusted for age, sex and/or site as covariates. Covariates were included in the analysis if statistically significant difference in the covariate was recorded between groups. Each donor is indicated by a unique symbol that is used consistently throughout all figures. Grey horizontal lines indicate the mean value derived from n = 9 Healthy donors. A description of the Healthy donor cohort is presented in Supplementary Table 3.

We next sought to assess if the increased trace-level of cytokines detected in the plasma of PASC-CVS donors could have physiological consequences. To assess this and given the cardiovascular symptoms reports by PASC-CVS participants, we cultured human induced pluripotent stem cell-derived cardiomyocytes (hiPSCs) and then added a cocktail of IL-1β, IL-6, IL-12 and MCP-1 (Figure 4). Cytokines were added at concentrations reflecting those detected in the Recovered cohorts (‘Recovered cytokine mimic’) or the PASC-CVS cohorts (‘PASC-CVS cytokine mimic’) (Figure 4). Cardiomyocyte function (as determined by amplitude, beat rate, upstroke velocity and relaxation velocity) was then assessed over the ensuing 48 hours (Figure 4). At 24- and 48-hours post-treatment PASC-CVS cytokine mimic treated samples had a reduced amplitude and reduced upstroke velocity compared to Recovered cytokine mimic treated and control samples (Figure 4). At 48 hours post-treatment PASC-CVS cytokine mimic treated samples had enhanced relaxation velocity compared to Recovered cytokine mimic treated and control samples (Figure 4). Interestingly, limited effects on cardiomyocyte function were observed when cardiomyocytes were treated with individual cytokines, suggesting that the combination of cytokines had a synergistic effect (Supplementary Figure 5).

**Figure 4:**
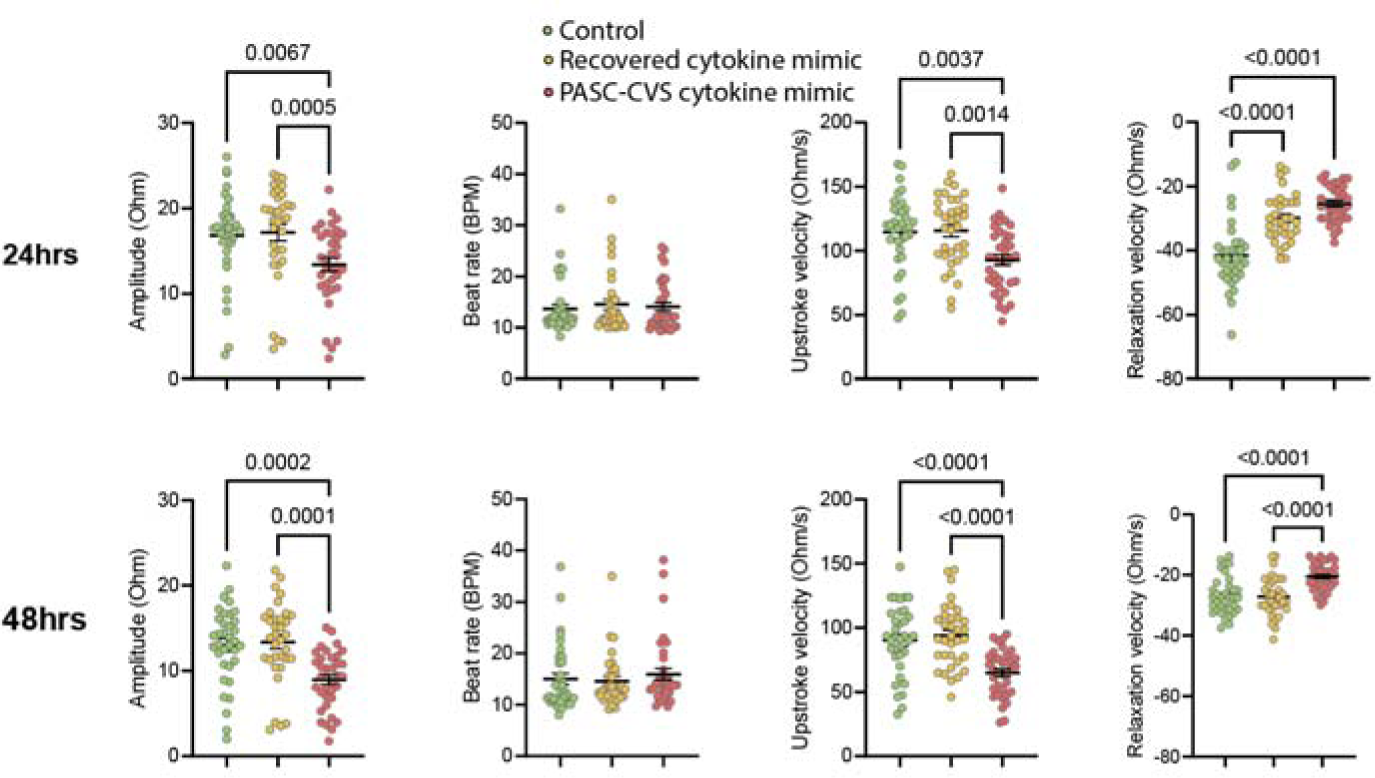
Trace-level pro-inflammatory cytokine cocktail affects cardiomyocyte function. Primary human cardiomyocytes were incubated for 24 hours (top line) and 48 hours (bottom line) a with media alone ((RPMI-1640/B27 with insulin; control) or a cocktail of IL-12, IL-1β, MCP-1 and IL-6 (‘cytokines’). Cytokine concentrations were either ‘Recovered cytokine mimic’ (IL-12: 0.007fg/mL + IL-1β 0.01fg/mL + MCP-1 0.01fg/mL + IL-6 0.004fg/mL) or ‘PASC-CVS cytokine mimic’ (IL-12: 41fg/mL + IL-1β 21fg/mL + MCP-1 14fg/mL + IL-6 21fg/mL) to reflect levels detected in the PASC-CVS cohorts. Graphs show mean ± SEM. Normal distribution of data was assessed with the Shapiro-Wilk test. Statistical significance was determined with a Kruskal-Wallis test with Dunn’s multiple comparison test or Welch ANOVA test and Dunnett’s multiple comparison test.

Given the effect of treating cardiomyocytes with cytokines, we hypothesised that the plasma of PASC-CVS donors would also differentially affect cardiomyocyte function relative to that of Recovered plasma. To test this hypothesis, cardiomyocytes were incubated for 48-hours with PASC-CVS donor plasma (Figure 5). PASC-CVS plasma treated cells had a significantly reduced relaxation velocity at 24-hours post-treatment and a reduced amplitude and upstroke velocity at 48-hours post-treatment compared to Recovered plasma treated cells (Figure 5). The effect of PASC-CVS plasma on cardiomyocyte upstroke velocity was not observed in the presence of dexamethasone at 24 hours post-treatment (Supplementary Figure 6). To further confirm the effects of donor plasma on cardiomyocyte function, RNAseq was performed on plasma treated cardiomyocytes at 24 hours post-treatment (Figure 6). Upregulated pathways in PASC-CVS plasma treated cardiomyocytes (relative to those treated with Recovered plasma) included ‘angiogenesis’, ‘inflammatory response’, ‘coagulation’, ‘hypoxia’ and ‘hypertrophic cardiomyopathy’ (Figure 6).

**Figure 5:**
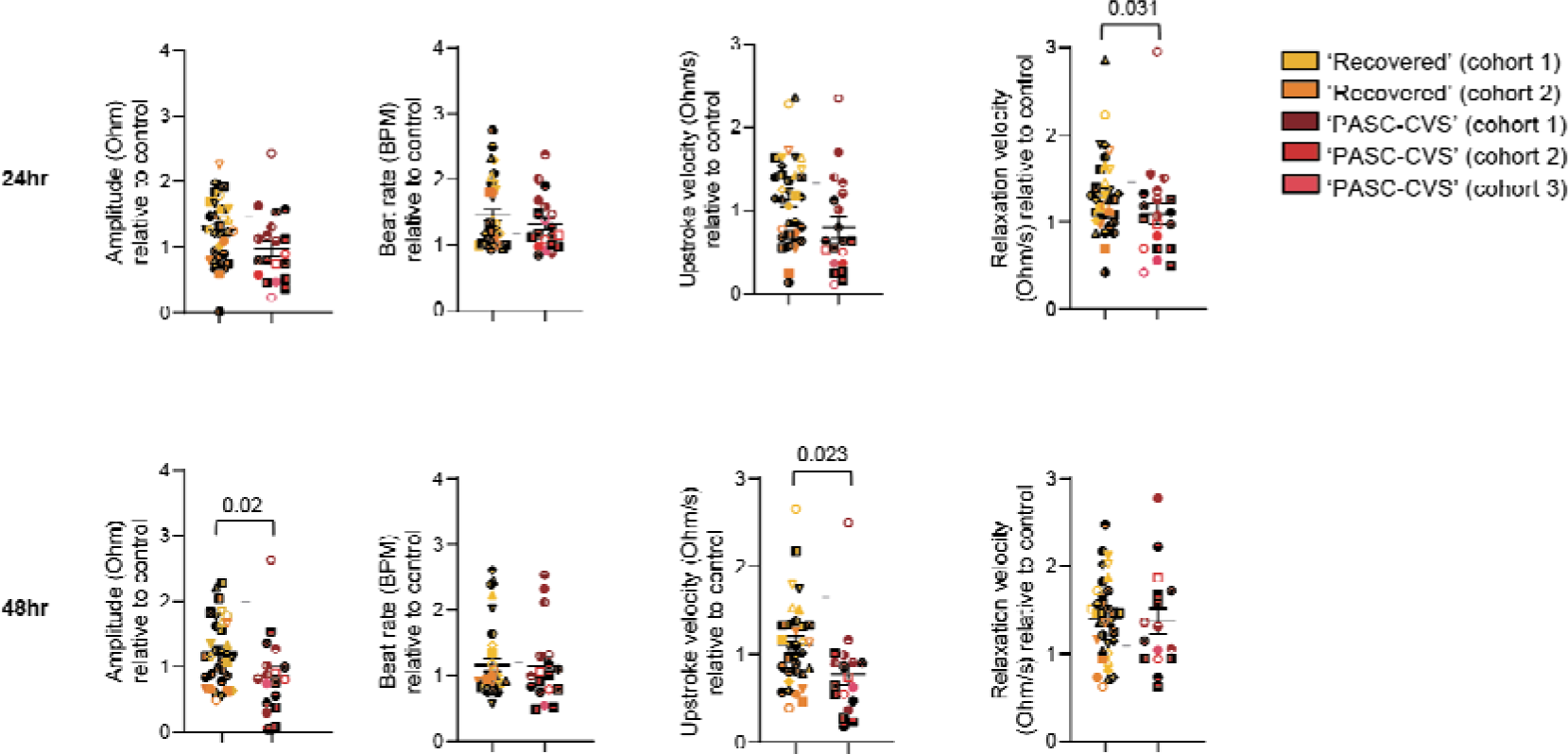
Reduced relaxation velocity, amplitude and upstroke velocity in cardiomyocytes after treatment with PASC-CVS plasma. Mean ± SEM is shown in all graphs. Statistical significance was determined with an ANCOVA adjusted for age, sex and/or site as covariates. Covariates were included in the analysis if statistically significant difference in the covariate was recorded between groups. Each donor is indicated by a unique symbol that is used consistently throughout all figures. Grey horizontal lines indicate the mean value derived from n = 22 Healthy donors. A description of the Healthy donor cohort is presented in Supplementary Table 3.

**Figure 6:**
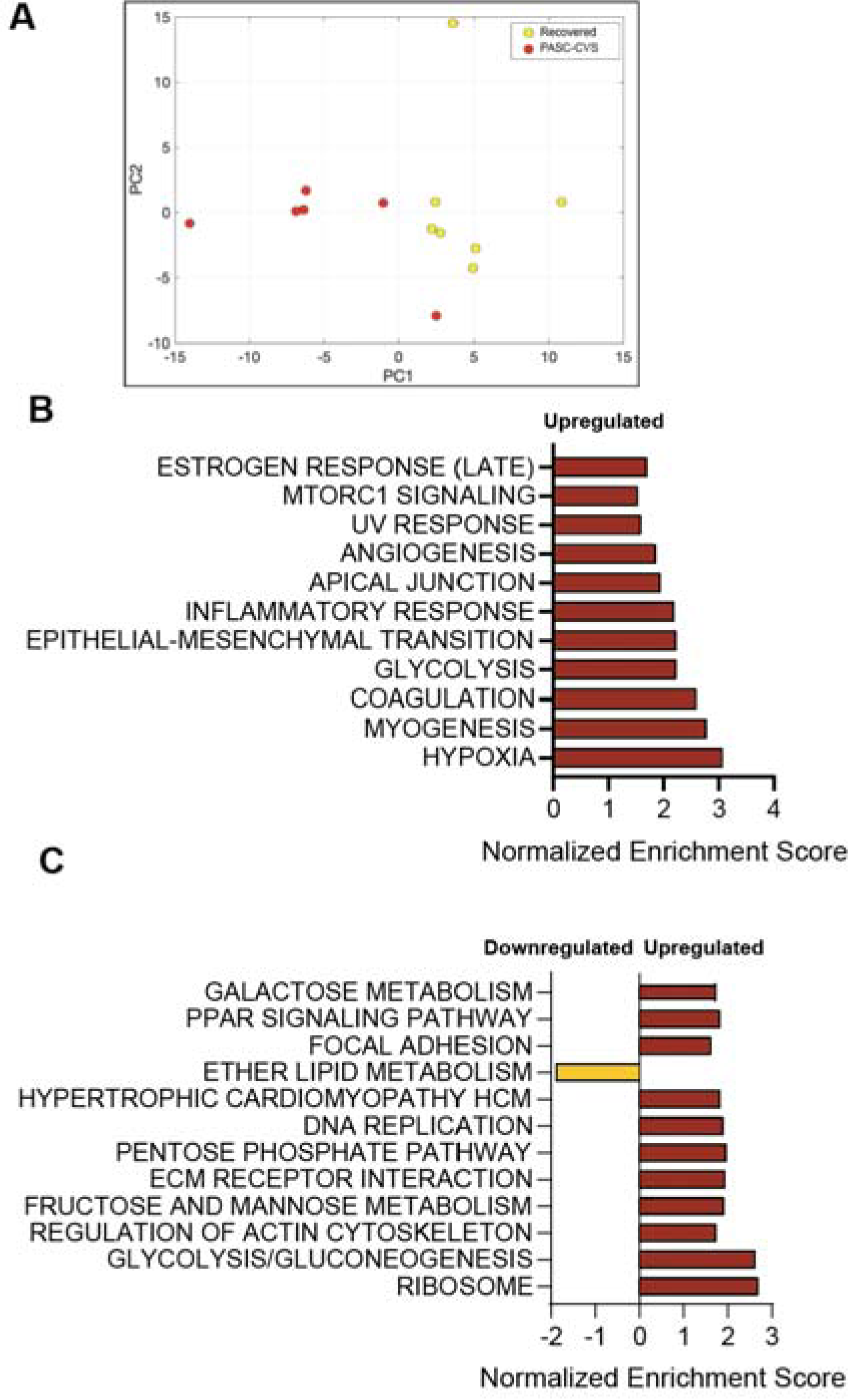
PASC-CVS plasma alters the transcriptome of cardiomyocytes relative to those treated with Recovered plasma. **(A)** PCA plot of cardiomyocyte transcripts following treatment with PASC-CVS or Recovered plasma for 24 hours **(B)** Pathway analysis of transcripts derived from PASC-CVS treated cardiomyocytes relative those treated with Recovered plasma analysed using the mSigDB Hallmark database. **(C)** Pathway analysis of transcripts derived from PASC-CVS treated cardiomyocytes relative those treated with Recovered plasma analysed using the mSigDB Hallmark database. **(B & C)** Only pathways with an adjusted p value of <0.05 are shown.

Additional differences in cardiomyocyte function were recorded following treatment with recombinant cytokines (Figure 4) as compared to treatment with donor plasma (Figure 5). This likely reflects the presence of other differentially abundant proteins in donor plasma. To explore this in more detail we performed proteomics on plasma from Recovered and PASC-CVS donors (Table 2). Many proteins were more abundant in plasma from PASC-CVS donors than from Recovered donors (Table 2; Supplementary Figure 7). Of particular interest, several proteins with increased abundance in PASC-CVS donors were associated with coagulation and platelet activation (e.g. Prothrombin, Serum amyloid A-4 protein and CXCL7) (Table 2; Supplementary Figure 6), and numerous proteins with increased abundance in PASC-CVS donors were associated with the complement cascade (e.g. Complement C1r subcomponent, Complement factor B and Complement component C9) (Table 2; Supplementary Figure 6). Proteins relating to coagulation and complement were also more abundant in PASC-CVS plasma compared to that of Healthy donors (Supplementary Table 4) whilst numerous proteins were of decreased abundance in PASC-CVS donors compared to Healthy donors (Supplementary Table 5). A finite set of proteins were more abundant in Recovered compared to Healthy donors (Supplementary Table 6), whilst a large number of proteins were less abundant in the plasma of Recovered donors compared to Healthy donors (Supplementary Table 7). The proteins that were less abundant in the plasma of PASC-CVS and Recovered compared to Healthy donors were mostly cytoplasmic and associated with vesicles (Supplementary Tables 5 and 7).

**Table 2:** Proteins with increased abundance in the plasma of PASC-CVS donors compared to Recovered donors.

| **Protein** | **UniProtKB accession** | **p-value<sup>1</sup>** |
| --- | --- | --- |
| Complement C1r subcomponent (EC 3.4.21.41) | P00736 | 0.0390 |
| Alpha-2-HS-glycoprotein (Alpha-2-Z-globulin) | P02765 | 0.0352 |
| Prothrombin (EC 3.4.21.5) | P00734 | 0.0130 |
| Inter-alpha-trypsin inhibitor heavy chain H4 | Q14624 | 0.0291 |
| Profilin-1 | P07737 | 0.0268<sup>^</sup> |
| Complement factor B (EC 3.4.21.47) | P00751 | 0.0197 |
| Complement component C9 | P02748 | 0.0035 |
| Alpha-1-acid glycoprotein 1 | P02763 | 0.0014 |
| Plasma kallikrein | P03952 | 0.0273 |
| Clusterin | P10909 | 0.0256 |
| Complement C1r subcomponent-like protein | Q9NZP8 | 0.0013 |
| Lysozyme C | P61626 | 0.0261 |
| Serum amyloid A-4 protein | P35542 | 0.0171 |
| Complement factor H | P08603 | 0.0042 |
| Inter-alpha-trypsin inhibitor heavy chain H3 | Q06033 | 0.0115 |
| C4b-binding protein alpha chain (C4bp) | P04003 | 0.0190 |
| Complement factor I | P05156 | 0.0273 |
| Alpha-1-acid glycoprotein 2 | P19652 | 0.0208 |
| Immunoglobulin lambda variable 3-10 | A0A075B6K4 | 0.0059<sup>^</sup> |
| Platelet basic protein (PBP) (C-X-C motif chemokine 7) | P02775 | 0.0366<sup>^</sup> |
| Immunoglobulin heavy variable 1-3 | A0A0C4DH29 | 0.0311 |

<sup>1</sup>. Determined by ANCOVA adjusting for age, sex and site
<sup>^</sup> Variance homogeneity or approx. equal sample size were evaluated for appropriateness of ANCOVA testing. These proteins did not meet this assumption.

Finally, we sought to identify if nanotechnology (referred to as the cardiac chip) could be used to identify trace-levels of biomarkers associated with cardiovascular disease, namely B-Type Natriuretic Peptide (BNP), Atrial natriuretic peptide (ANP), MIP-1β, cardiac troponin I (cTnI) and Creatine Kinase MB (CK-MB) (Figure 7). There were significantly lower levels of MIP-1β detected in PASC-CVS donors compared to Recovered individuals (Figure 7). Interestingly, there was a trend towards elevated MIP-1β, cTnI and CK-MB levels in both Recovered and PASC-CVS donors compared to the mean of Healthy donors (Figure 7).

**Figure 7:**
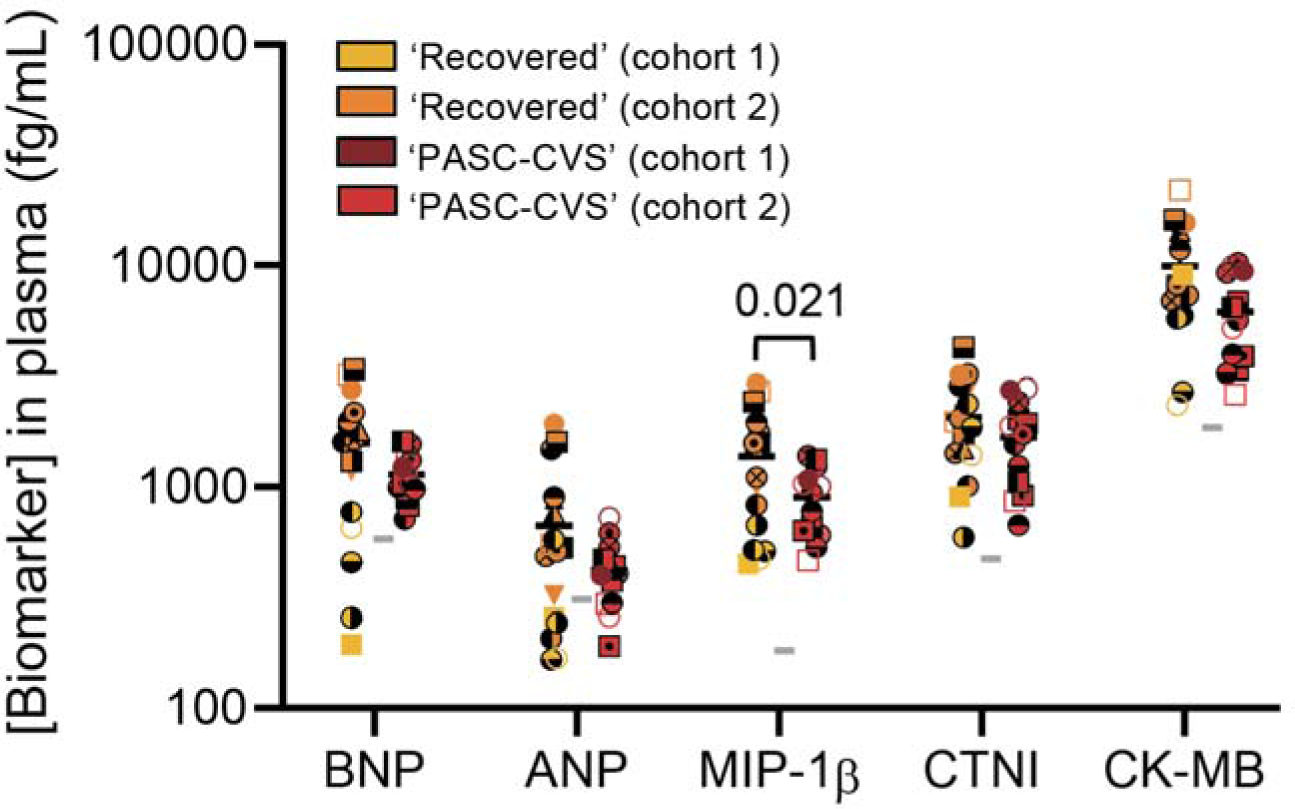
No increase in cardiovascular damage markers in PASC-CVS cohorts. Levels in plasma were determined using the cardiac chip. Statistical significance was determined with an ANCOVA adjusted for age, sex and/or site as covariates. Covariates were included in the analysis if statistically significant difference in the covariate was recorded between groups. Each donor is indicated by a unique symbol that is used consistently throughout all figures. Grey horizontal lines indicate the mean value derived from n = 22 Healthy donors. A description of the Healthy donor cohort is presented in Supplementary Table 3.

## DISCUSSION

This study demonstrates that PASC-CVS donors have a distinct pro-inflammatory blood transcriptome >1 year post-SARS-CoV-2 infection. PASC-CVS donors also have elevated levels of trace-level plasma cytokines at approximately 18-months post-infection which are associated with the altered functionality of human pluripotent stem cell derived cardiomyocytes *in vitro*. Proteomics demonstrated further differences in PASC-CVS and Recovered plasma including enrichment of complement and coagulation associated proteins.

The presence of elevated cytokines in the plasma of PASC-CVS donors is consistent with previous descriptions that eight to ten months post-infection patients with PASC have low level elevations in proinflammatory cytokine levels^13-15^. However, to the best of our knowledge the present study represents the longest period between SARS-CoV-2 diagnosis and sample collection (including samples collected >19 months post-infection) to show elevated cytokine levels in the plasma of participants with PASC. The present study was also the first to specifically identify elevated trace-level cytokines in individuals living with PASC-CVS. We speculate that the prolonged period post-infection of the present study is what necessitates the use of ultra-sensitive nanotechnology, as cytokines detected by conventional methods (e.g. a bead-based multiplex assay or ELISA) in our donor samples were often below the detection limit of the assay. These data raise the exciting possibility that nanotechnology could be used to improve the diagnosis of people living with PASC-CVS, in particular those who have been living with this condition for more than one year. Accordingly, the sensitivity and specificity of the immunostorm chip for PASC-CVS diagnosis requires further research with appropriately powered cohorts. Importantly, while the current work was performed using an advanced confocal Raman microscope, transition to more clinically applicable instrumentation such as fluorescence microscopy and microfluidics is highly feasible and likely to reduce costs and complexity^39,40^.

The presence of inflammatory cytokines is known to affect cardiovascular function. For example, a cocktail of IFNγ, IL-1β and poly(I:C) induced diastolic dysfunction in human cardiac organoids^41^. However, it is important to note that such studies have investigated the effects of cytokines in acute COVID-19, which are typically within the picogram to nanogram range^41^. Here, we observed a clear effect of cytokines in the femtogram range, suggesting that the trace-level cytokines in PASC-CVS individuals have physiological consequences. The mechanisms by which cytokines depress cardiac function has been studied extensively with promiscuous effects reported across diverse mechanisms of cardiac signalling and excitation-contraction coupling^42^. Clinical studies evaluating concentration-dependent effects of cytokines in heart failure patients have shown a significant correlation between circulating cytokine levels and degree of heart failure, including contractile measures of ejection fraction^43^. This is in alignment with our study showing a cocktail of trace-level proinflammatory cytokines decreased the amplitude of cardiomyocyte contraction, a measure of pump performance. Differential gene expression analysis further validated the perturbations observed in cardiovascular function in PASC-CVS plasma treated cardiomyocytes. Differentially expressed pathways in PASC-CVS plasma treated cardiomyocytes included those associated with hypertrophic cardiomyopathy and the inflammatory response. Importantly, the effects of PASC-CVS plasma on cardiomyocyte upstroke velocity were not observed in the presence of the anti-inflammatory dexamethasone. The potential of nanotechnology to identify individuals with PASC-CVS with chronic inflammation, such that they could be administered the appropriate anti-inflammatories to reduce cardiovascular symptoms is exciting.

With the exception of CXCL7 no other cytokine was detected as enriched in PASC-CVS plasma using proteomics. This is consistent with trace-level nature of the detected cytokines and the detection limit of proteomics in the absence of an enrichment step^44^. However, whole plasma proteomics did reveal several other important differences between Recovered and PASC-CVS samples. Firstly, several proteins associated with coagulation and clotting were enriched in PASC-CVS samples. For example, Serum amyloid A4 protein was enriched in PASC-CVS samples and this protein is known to enhance clotting in plasma and increase the risk of venous thromboembolism^45^. Interestingly, previous studies have described large anomalous (amyloid) deposits in the plasma of PASC patients, which consist primarily of α(2)-antiplasmin, various fibrinogen chains as well as Serum Amyloid A4^46^. Our data are thus consistent with a microclotting phenotype in PASC-CVS^47^. A second key feature of these proteomic data is the elevated level of proteins associated with the complement cascade in the plasma of PASC-CVS donors. These findings are consistent with recent studies describing elevated levels of classical, alternative and/or terminal pathways of complement activation in patients with PASC^5,48,49^. It is also possible that these proteins represent an alternative diagnostic target for PASC. Indeed, patient age, C5bC6/C7 complex ratio, vWF/ADAMTS13 ratio and patient body mass index were sufficient to predict 12-month post-infection PASC (mean area under the curve of 0.81 ± 0.08)^49^. Others have reported that the activation fragments iC3b, terminal complement complex, Ba, and C5a had a receiver operating characteristic predictive power of 0.785 in predicting PASC^48^. Given the independent verification of complement dysregulation in PASC-CVS provided by the current study the diagnostic potential of various components of the complement cascade requires further inquiry.

The question remains as to why individuals with PASC-CVS have chronic low-level inflammation as well as signs of coagulation and complement activation. It is possible that these phenomena are not mutually exclusive. For example, Cervia-Hasler and colleagues suggested increased membrane insertion of terminal complement complexes contributed to tissue damage and a thromboinflammatory signature in PASC patients and that increased cleavage by thrombin further drove complement activation^49^. Indeed, it is possible that complement activation drives not only tissue damage but also the trace-level cytokine signature observed in the present study^50,51^. Should this be the case, the question remains as to what the underlying trigger for complement activation (and the resultant inflammatory signature) is in PASC patients. Cervia-Hasler and colleagues suggest that this may be the result of antibody mediated activation of the classical complement pathway as the result of reactivation of viruses such as cytomegalovirus or Epstein-Barr virus^49^. Alternatively, it is possible that complement activation and inflammation is triggered by the persistence of SARS-CoV-2-derived pathogen associated molecular patterns (PAMPs). Dissection of these different mechanisms of PASC pathogenesis has important implications for patient care. For example, an understanding of the prevalence of viral reactivation in PASC patients will inform the need for antiviral treatment in these individuals.

Traditional markers of cardiovascular disease (e.g. CK-MB and cTnI), even when measured by nanotechnology, were not significantly different between PASC-CVS and Recovered donors. This adds further weight to the potential of using trace-level cytokines, rather than more traditional markers of cardiovascular disease, for the diagnosis of PASC-CVS. Interestingly, MIP-1β was significantly lower in PASC-CVS individuals compared to those who recovered from SARS-CoV-2. This speaks to a level of specificity in the elevated cytokine/chemokine signal in PASC-CVS. When compared to the mean value of Healthy donors (i.e. those without a SARS-CoV-2 infection) it was striking to notice a pattern of elevated levels of MIP-1β, CK-MB and cTnI in both the Recovered and PASC-CVS samples. cTnI is a well-established sensitive and specific marker of myocardial infarction^52^. Elevated CK-MB can also be used for the diagnosis of myocardial infarction^53,54^. It is also interesting to note the pattern of altered plasma proteome, with reduced abundance of vesicle-associated proteins in the plasma of both Recovered and PASC-CVS donors compared with Healthy donors. Whole blood transcriptomics further identified numerous differentially expressed genes between Recovered individuals and Healthy Controls. These data are all potentially consistent with prior studies demonstrating that SARS-CoV-2 infection can induce numerous long-term changes, including those associated with accelerated aging^55,56^. Indeed, in the context of cardiovascular disease, Poyatos and colleagues showed that post-COVID-19 patients still displayed elevated troponin and NT-proBNP levels up to 12-months post-infection. However, the comparison to Healthy controls in the present study must be interpreted with caution. It is important to recognise that in the cytokine and proteomic analysis the Healthy samples were a historical cohort and therefore must be considered as a separate site to the more recently acquired Recovered and PASC-CVS samples. Accordingly, site and donor group both form mutually exclusive events for the Healthy controls and therefore any ANCOVA analysis with the Healthy samples cannot control for site as a covariate. Accordingly, here we have elected to limit comparisons with the Healthy cohort in order to interpret the data in a conservative manner. However, these data do represent an impetus for further investigation as to the effects of SARS-CoV-2 infection on the cardiovascular system, even in individuals who do not experience prolonged symptoms of viral infection.

There are several important limitations of the present study to note. Firstly, the focus on individuals with prolonged PASC-CVS limited the sample number available for the present study. Accordingly, these studies require validation in additional patient cohorts including more contemporary cohorts who were infected with more recent SARS-CoV-2 strains. It would also be important to investigate whether the observations described herein applied to other subtypes of PASC (e.g. neurological disease, respiratory disease etc.). Indeed, individuals with PASC-CVS were recruited to the present study based on the presence of chest pain and/or heart palpitations. Perhaps not surprisingly, given the multifaceted impacts of cardiovascular disease, numerous individuals reported other concurrent symptoms such as shortness of breath and fatigue. Whether the observed phenotype would apply to individuals with exclusively chest pain and/or heart palpitations (of which there were only two in the present study) remains to be determined. However, despite these limitations this study provides important new insights into the complex disease that is PASC and offers opportunities to improve the diagnosis, treatment and understanding of the ever-growing number of individuals living with PASC-CVS.

## ACKNOWLEDGEMENTS

We thank all the participants for making this study possible. We also acknowledge Sam Eiszele and UQ Human Studies Unit. We thank research staff at the Royal Adelaide Hospital Intensive Care and Infectious Diseases Units (Adelaide, SA) for participant recruitment, sampling and follow up. The authors appreciate receiving the technical and scientific guidance from the Queensland node of the Australian National Fabrication Facility (Q-ANFF) in confocal Raman mapping, Dr Elliot Cheng from Centre for Microscopy and Microanalysis (CMM, University of Queensland) in nanofabrication, and Dr Amanda Nouwens from the Mass Spectrometry Facility of the School of Chemistry and Molecular Biosciences, The University of Queensland.

