## Supplementary Figures for "Cardiovascular symptoms of PASC are associated with trace-level cytokines that affect the function of human pluripotent stem cell derived cardiomyocytes"

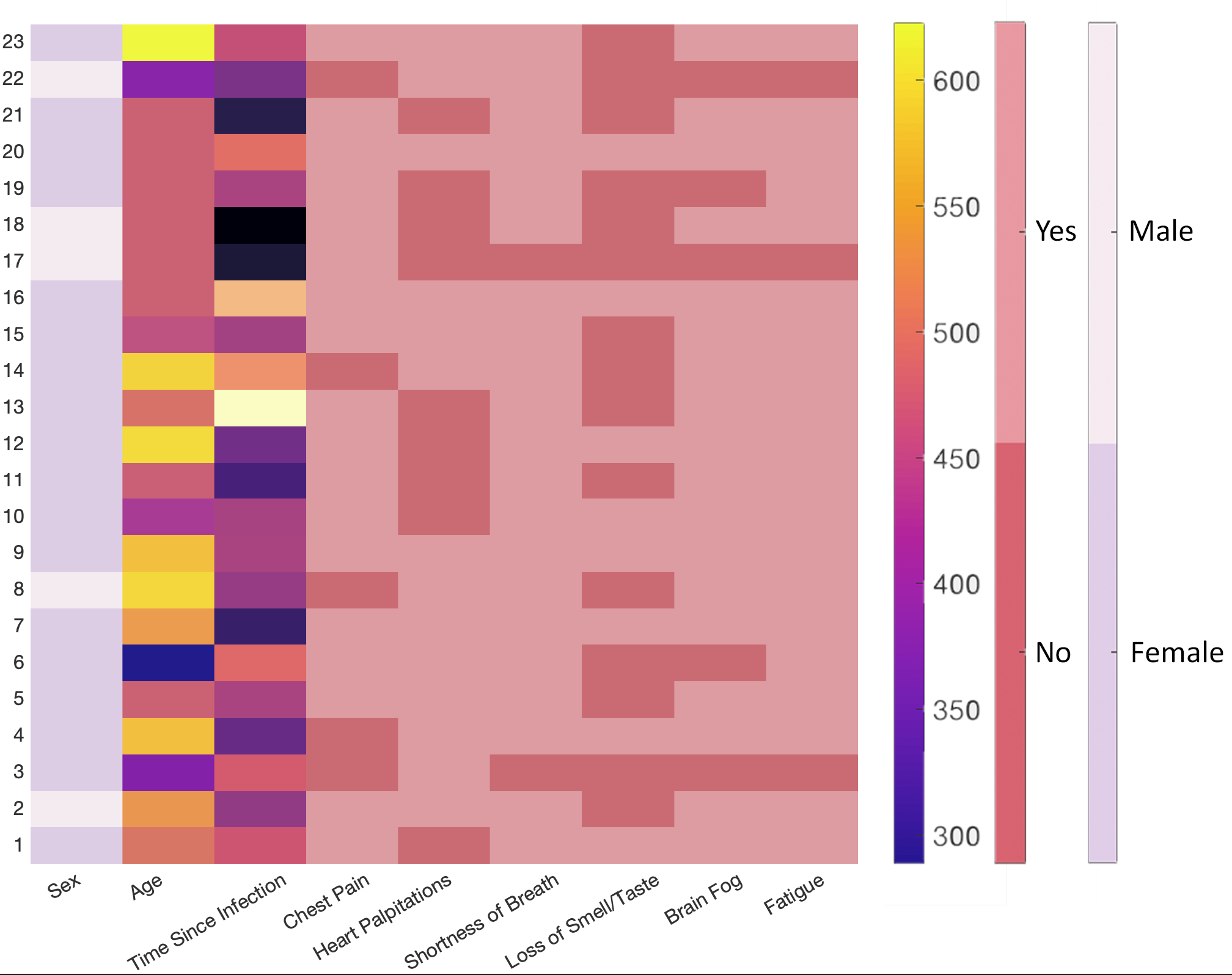

**Supplementary Figure 1: Symptoms described by the 23 PASC-CVS participants.**

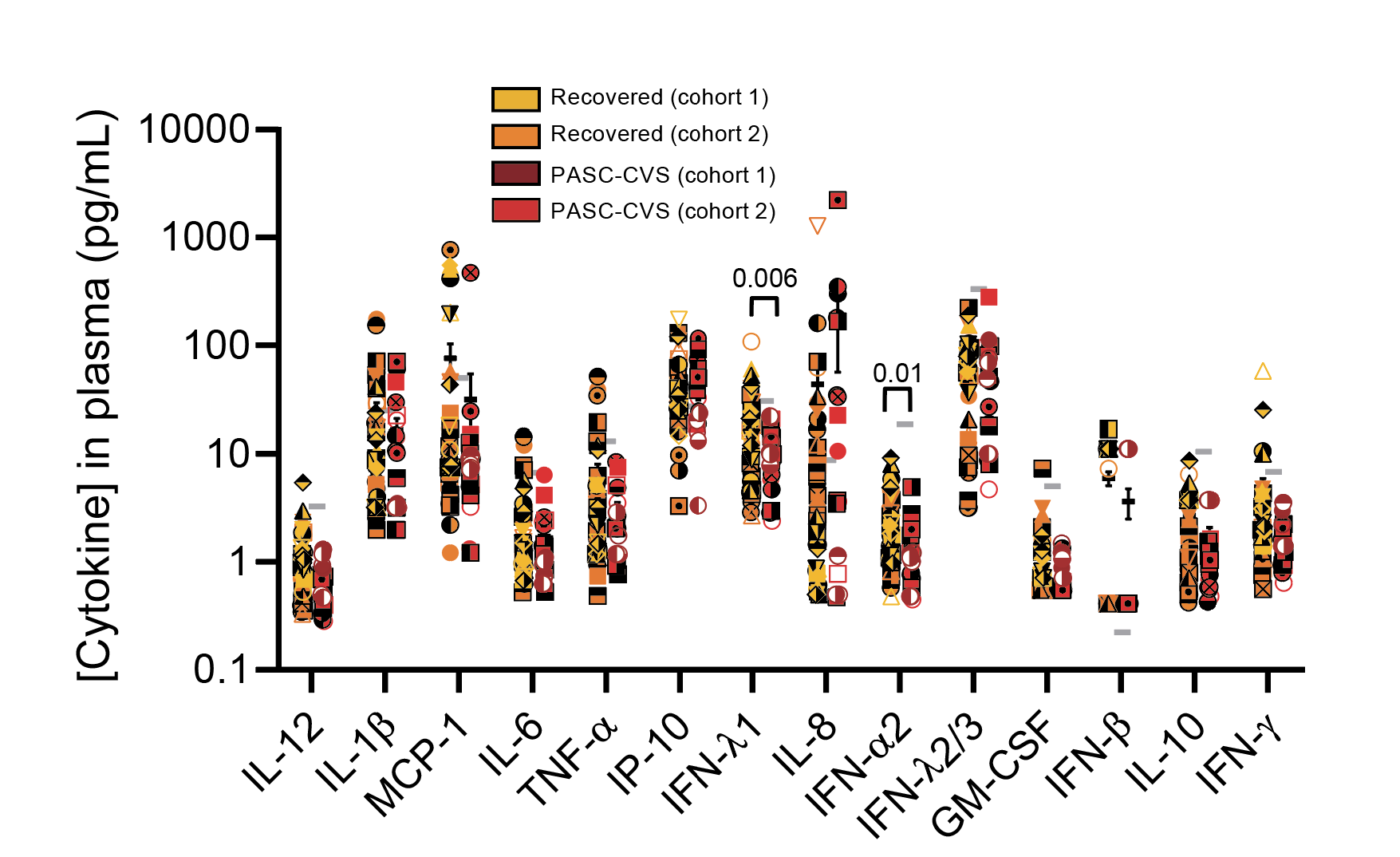

**Supplementary Figure 2**: **Cytokine levels in the plasma of recovered and PASC-CVS donors**. Cytokine levels were detected in the plasma of participants using an ELISA (MCP-1) or a Legendplex bead-based assay (all other cytokines). Statistical significance was determined with an ANCOVA adjusted for age, sex and/or site as covariates. Covariates were included in the analysis if statistically significant difference in the covariate was recorded between groups. Values below the detection limit of the cytokine in question were taken as the assay lower bound. Each donor is indicated by a unique symbol that is used consistently throughout all figures. Mean ± SEM is shown. Grey horizontal lines indicate the mean value derived from n = 16 Healthy donors. A description of the Healthy donor cohort is presented in Supplementary Table 6.

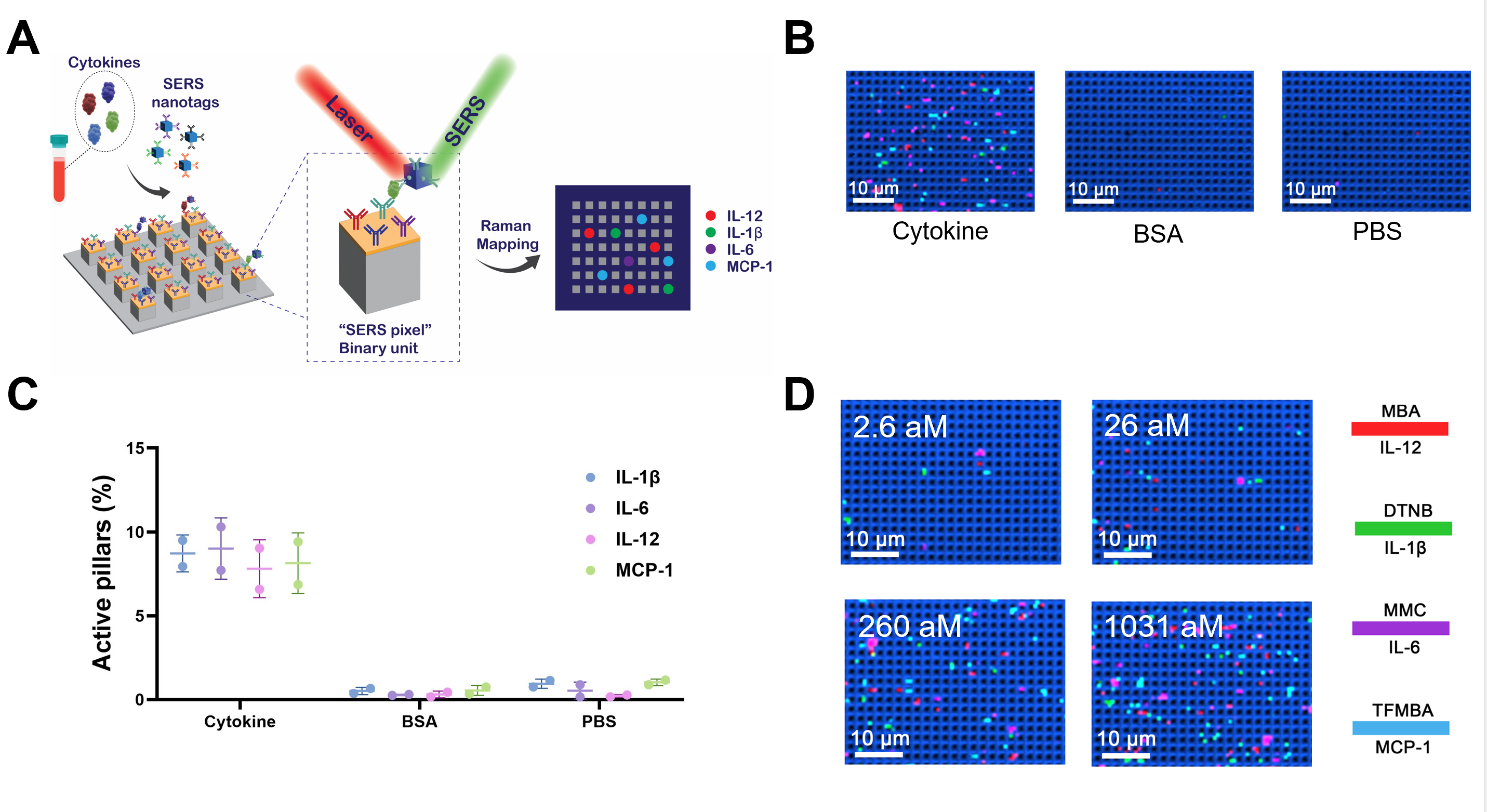

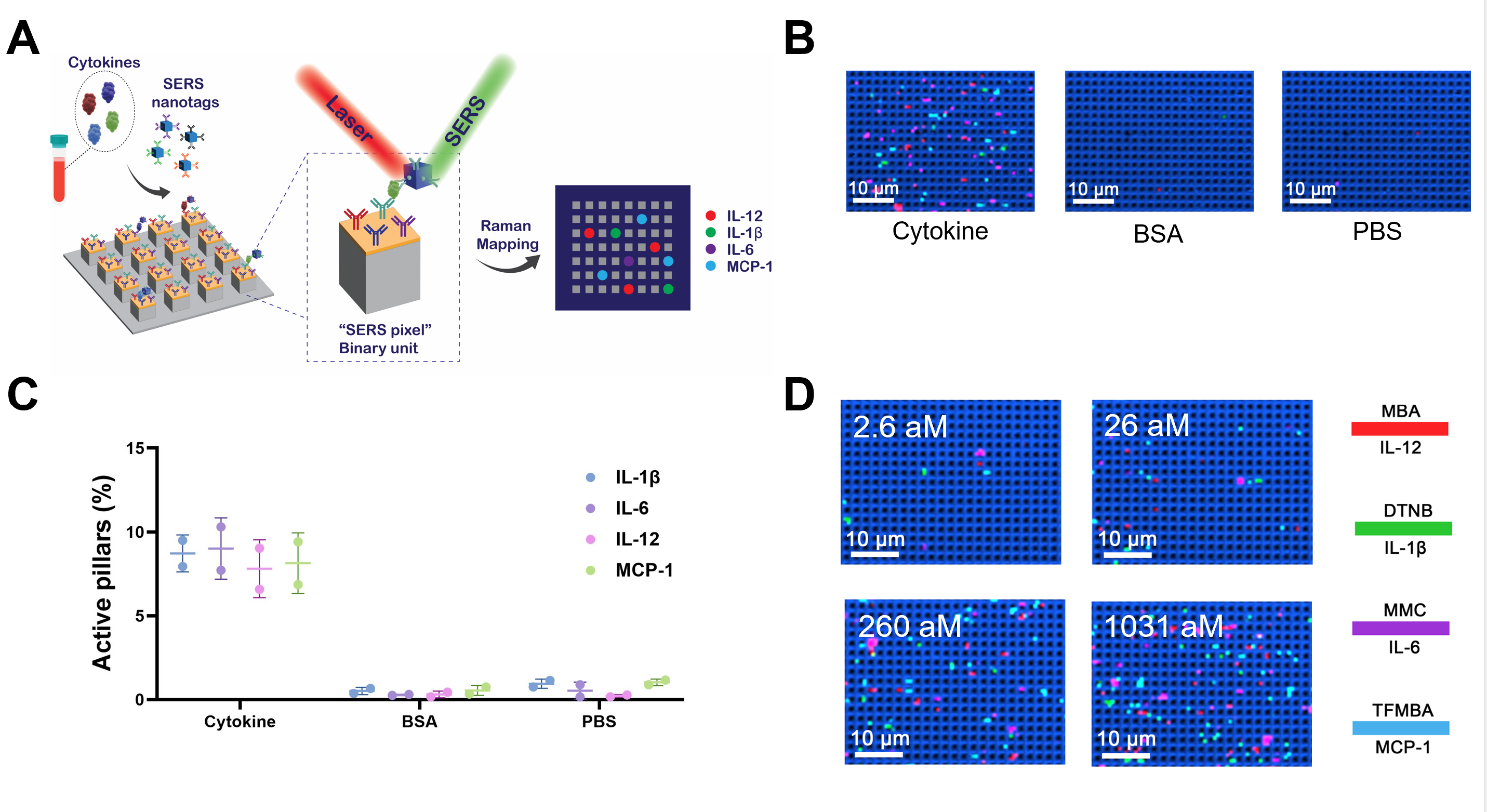

**Supplementary Figure 3: The use of the immunostorm chip to detect trace-level cytokines in patient samples. A**) Schematic workflow: Cytokines are captured on a nanopillar array and labelled with single-particle active SERS nanotags to from an immune complex (‘SERS pixel'). Trace-level cytokine measurement is achieved by mapping the nanopillar array by confocal Raman spectroscopy to count the SERS pixels. (**B**) Representative false-colour Raman images following treatment with a cytokine cocktail, bovine serum albumin (BSA) or PBS and (**C**) Representative false-colour Raman images of samples containing cytokines with a concentration of 2.6 aM, 26 aM, 260 aM, and 1031 aM.

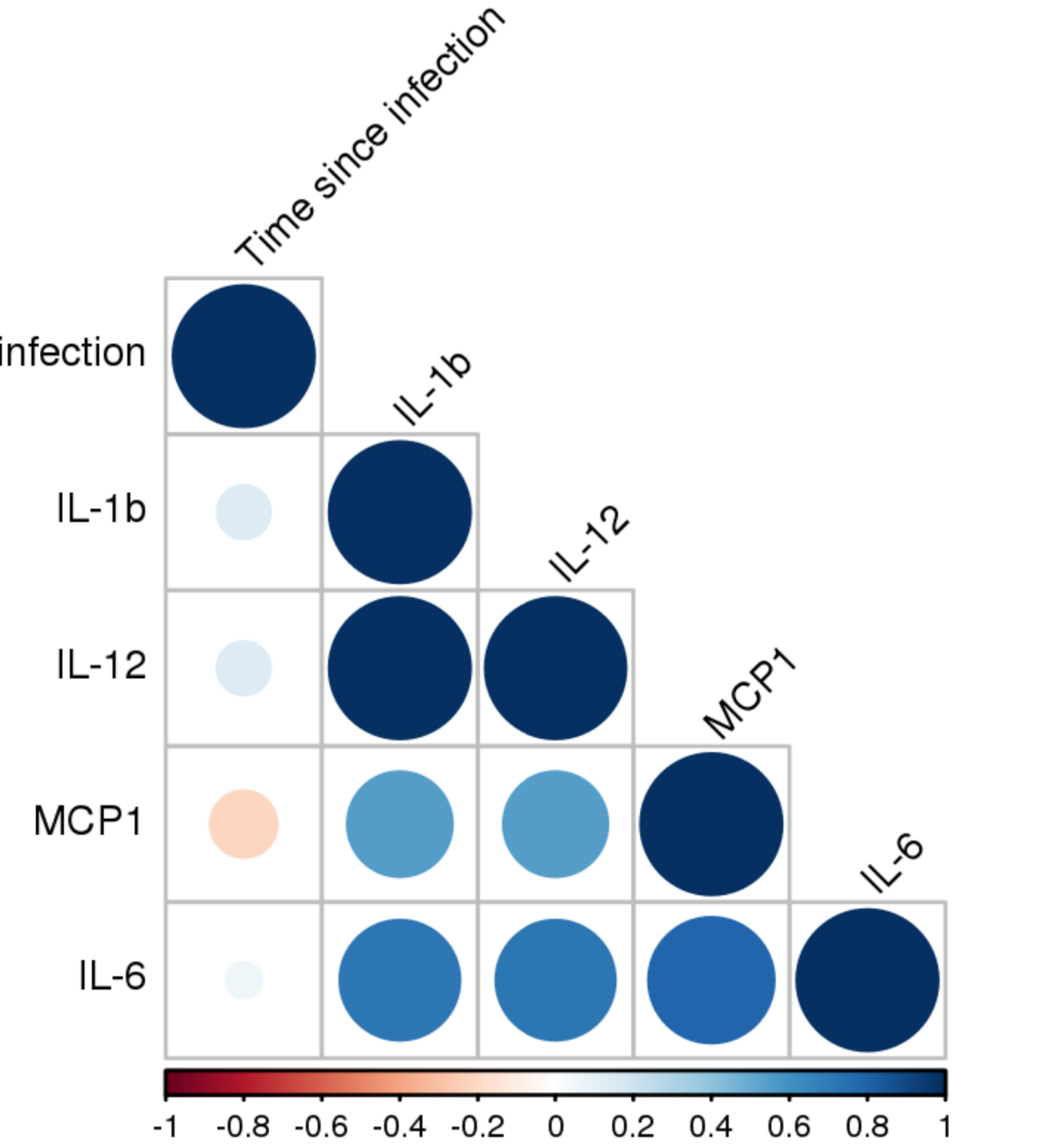

Time since infection

**Supplementary Figure 4: Pearson correlation matrix of trace-level cytokines detected by immunostorm chip in the plasma of PASC-CVS donors.**

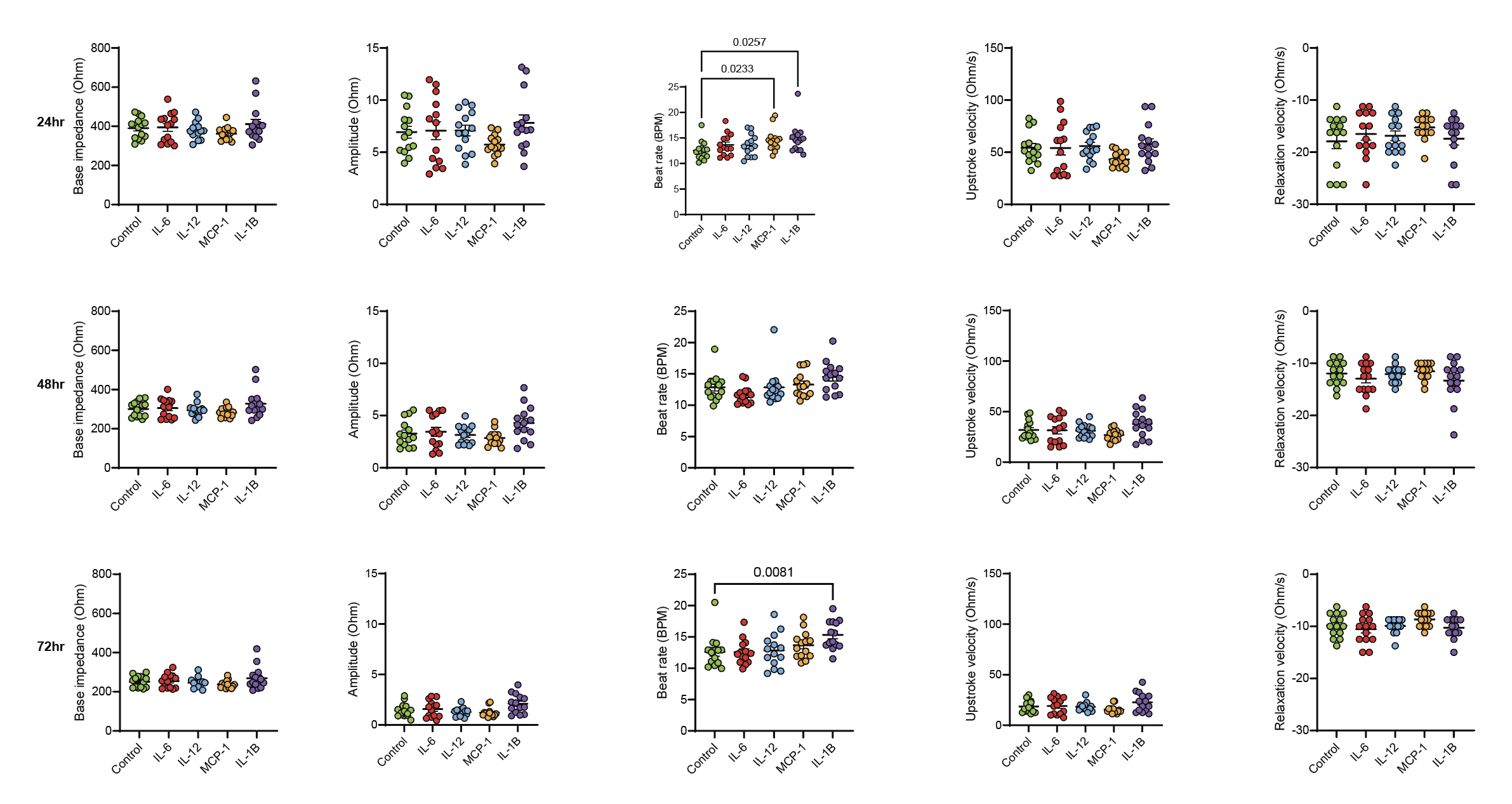

**Supplementary Figure 5:** **Trace-level individual pro-inflammatory cytokines did not affect cardiomyocyte function.** Human cardiomyocytes were incubated for 24 hours (top line) or 48 hours (bottom line) with media alone ((RPMI-1640/B27 with insulin; control) or individual pro-inflammatory cytokines. Cytokine concentrations were ‘PASC-CVS cytokine mimic’ (IL-12: 41fg/mL; IL-1β 21fg/mL; MCP-1 14fg/mL or IL-6 21fg/mL). Graphs show mean ± SEM. Normal distribution of data was assessed with the Shapiro-Wilk test. Mean ± SEM is shown. Statistical significance was determined with a Kruskal-Wallis test with Dunn’s multiple comparison test or Welch ANOVA test and Dunnett’s multiple comparison test.

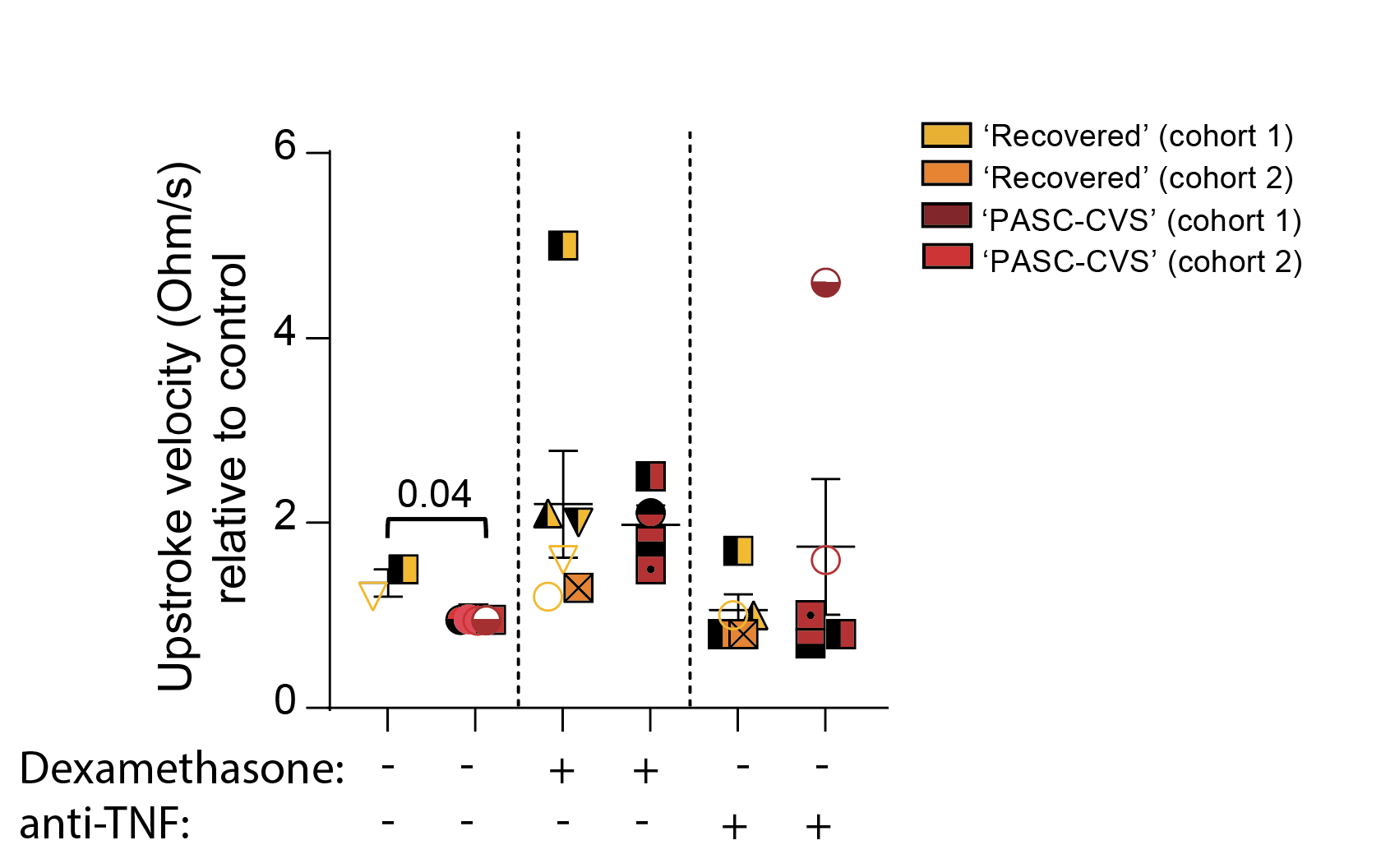

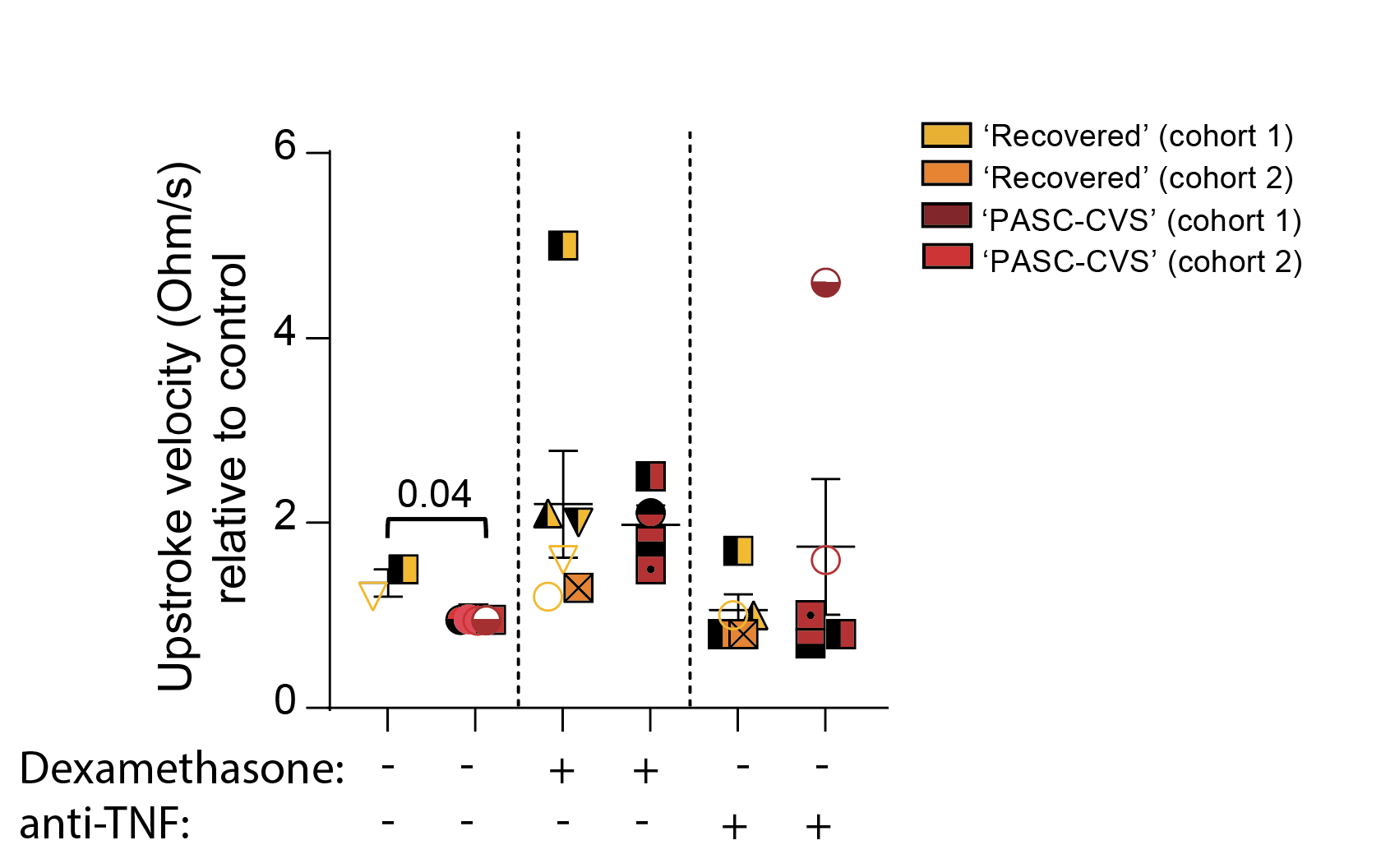

**Supplementary Figure 6: PASC-CVS plasma does not affect cardiomyocyte upstroke velocity in the presence of dexamethasone**. Primary human cardiomyocytes were incubated for 48 hours with age, sex and site matched PASC-CVS and Recovered plasma. Plasma was added in the presence or absence 100ng/mL of dexamethasone. Graphs sh	ow mean ± SEM. Normal distribution of data was assessed with the Shapiro-Wilk test. Mean ± SEM is shown. Statistical significance was determined with a Mann-Whitney U Test.

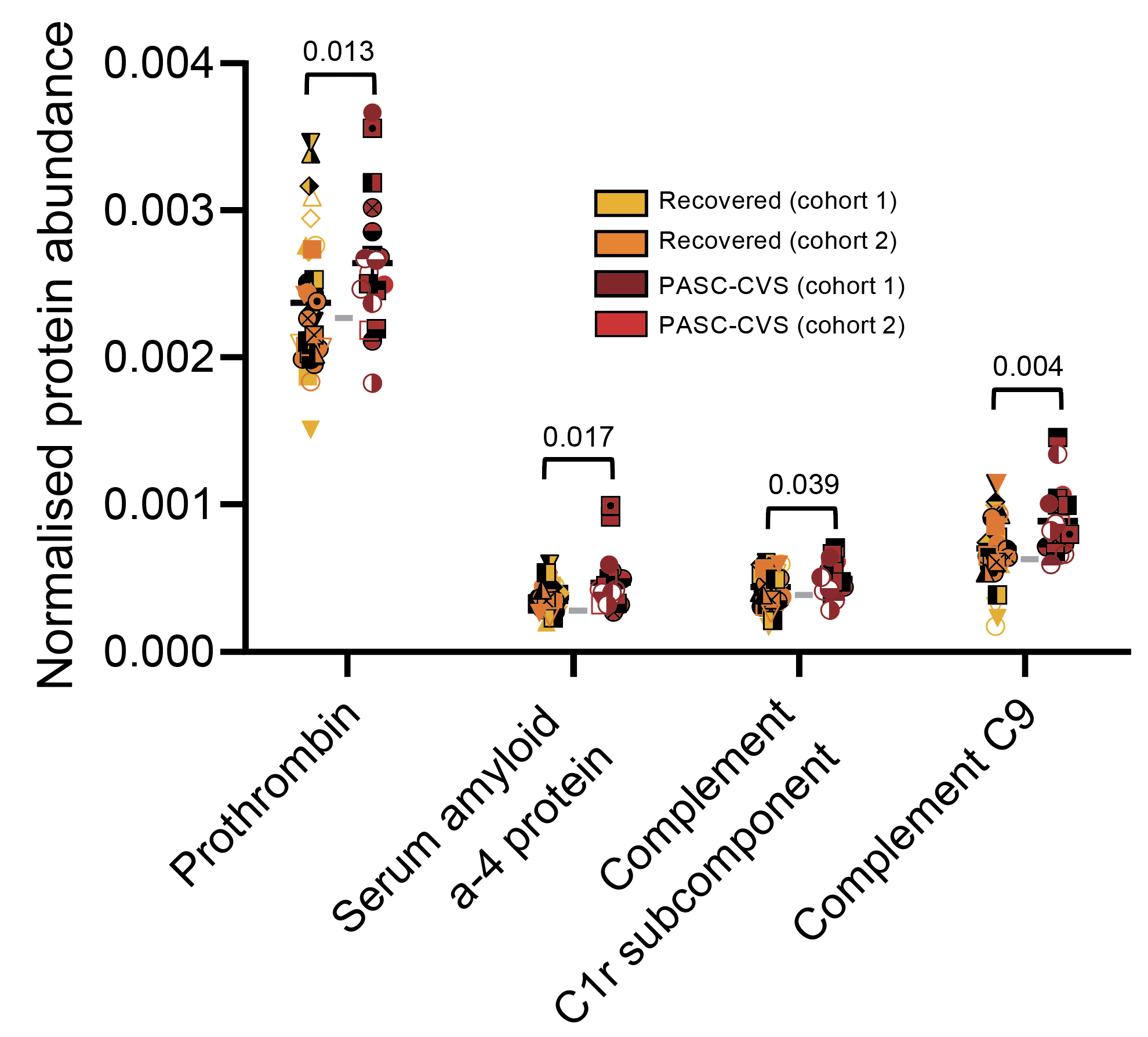

**Supplementary Figure 7: Normalised protein abundance in PASC-CVS and Recovered plasma.** Statistical significance was determined with an ANCOVA adjusted for age, sex and/or site as covariates. Covariates were included in the analysis if statistically significant difference in the covariate was recorded between groups. Each donor is indicated by a unique symbol that is used consistently throughout all figures. Mean ± SEM is shown Grey horizontal lines indicate the mean value derived from n = 22 Healthy donors. A description of the Healthy donor cohort is presented in Supplementary Table 6.
