## Supplementary Tables for "Cardiovascular symptoms of PASC are associated with trace-level cytokines that affect the function of human pluripotent stem cell derived cardiomyocytes"

**Supplementary Table 1: Participant characteristics in Cohort 1^1^**

|  | **Recovered** | **PASC** | **PASC-CVS** | **Healthy** | **P-value** |
| --- | --- | --- | --- | --- | --- |
| Total | 11 | 21 | 4 | 14 |  |
| Sex (M;F) | 5;6 | 13;8 | 1;3 | 7;7 | 0.531^2^ |
| Age (mean ± SEM) | 57.82(±3.3) | 52.29(±4.150) | 44.75(±9.286) | 50.5 (±3.905) | 0.5136^3^ |
| *Severity of acute infection^4^* |  |  |  |  | 0.18 |
| Mild | 9 | 11 | 3 | NA |  |
| Moderate | 2 | 2 | 0 | NA |  |
| Severe | 0 | 8 | 1 | NA |  |

^1.^Samples were obtained at 44 weeks (~308 days) and 68 weeks (~476 days) post-infection

^2.^Chi-squared test

^3^Kruskal-Wallis test

**Supplementary Table 2: Characteristics of participant plasma used for transcriptomic analysis of Cohort 2**

|  | **Recovered** | **PASC-CVS** | **P-value** |
| --- | --- | --- | --- |
| Total | 4 | 5 | - |
| Mean days post-infection that a blood sample was acquired (mean ± SEM) | 524.3±128.7 | 527.2±27.7 | 0.96^1^ |
| Sex (M;F) | 2;2 | 1;4 | 0.53^2^ |
| Age (mean ± SEM) | 52.52±21.46 | 45.85±3.96 | 0.51^1^ |
| *Severity of acute infection* |  |  | 0.21^2^ |
| Mild | 3 | 1 | - |
| Moderate | 1 | 4 |  |
| Severe | 0 | 0 |  |

1. Student’s t-test 2. Fisher’s exact test

**Supplementary Table 3 - Participant characteristics of the Healthy donor cohort 2**

|  | **Healthy** |
| --- | --- |
| Total | 22 |
| Age (mean ± SD) | 31 ± 13.8 |
| Sex (M/F) | 4/18 |
| Site |  |
| *Red Cross* | 9 |
| *Mater* | 13 |

**Supplementary Table 4 –Proteins with increased abundance in the plasma of PASC-CVS donors compared to Healthy donors**

| **Protein** | **UniProtKB accession** | **p-value^1^** |
| --- | --- | --- |
| Alpha-1-antitrypsin (Alpha-1 protease inhibitor) | P01009 | **0.0184** |
| Complement factor B (EC 3.4.21.47) | P00751 | **0.0188** |
| Complement factor H-related protein 2 | P36980 | **0.0267** |
| Complement component C9 | P02748 | **0.0086** |
| Alpha-1-acid glycoprotein 1 (AGP 1) | P02763 | **0.0101** |
| Apolipoprotein C-I | P02654 | **0.0014^** |
| Phosphatidylcholine-sterol acyltransferase | P04180 | **0.0274** |
| Serum amyloid A-4 protein | P35542 | **0.0006^** |
| Fibronectin | P02751 | **0.0333^** |
| Antithrombin-III | P01008 | **0.0025** |
| Complement factor H | P08603 | **0.0007** |
| Complement factor I | P05156 | **0.0029** |
| Alpha-1-acid glycoprotein 2 | P19652 | **0.0469** |
| Immunoglobulin heavy variable 1-46 | P01743 | **0.0099** |
| Apolipoprotein B-100 | P04114 | **0.0037^** |

**^1.^** Determined by ANCOVA adjusting for age and sex

**^** Variance homogeneity or approx. equal sample size were evaluated for appropriateness of ANCOVA testing. These proteins did not meet this assumption.

**Supplementary Table 5 –Proteins with decreased abundance in the plasma of PASC-CVS donors compared to Healthy donors**

| **Protein** | **UniProtKB accession** | **p-value^1^** |
| --- | --- | --- |
| Profilin-1 | P07737 | **0.0003** |
| Immunoglobulin heavy constant gamma 2 | P01859 | **0.0147** |
| Neutrophil defensin 1 | P59665 | **0.0248^** |
| Solute carrier family 2 | P11169 | **0.0469** |
| SUMO-specific isopeptidase USPL1 | Q5W0Q7 | **0.0362^** |
| Doublesex- and mab-3-related transcription factor 2 | Q9Y5R5 | **0.047** |
| Pyruvate kinase PKM | P14618 | **0.0015^** |
| Glyceraldehyde-3-phosphate dehydrogenase | P04406 | **0.0026** |
| Cofilin-1 | P23528 | **0^** |
| Beta-actin-like protein 2 | Q562R1 | **0.0062** |
| L-lactate dehydrogenase B chain | P07195 | **0.001^** |
| Transgelin-2 | P37802 | **0.0001^** |
| 14-3-3 protein zeta/delta | P63104 | **0.0002^** |
| Ras-related protein Rap-1b | P61224 | **0^** |
| Talin-1 | Q9Y490 | **0.0123^** |
| Vinculin (Metavinculin) | P18206 | **0^** |
| Integrin-linked protein kinase | Q13418 | **0.0037** |
| Fermitin family homolog 3 | Q86UX7 | **0.0003^** |
| Alpha-actinin-1 | P12814 | **0.0002^** |
| Ras-related protein Rab-15 | P59190 | **0.0024** |
| SH3 domain-binding glutamic acid-rich-like protein 3 | Q9H299 | **0.0002** |
| Peptidyl-prolyl cis-trans isomerase A | P62937 | **0.0002^** |
| Triosephosphate isomerase | P60174 | **0.0338** |
| Fructose-bisphosphate aldolase A | P04075 | **0.0123** |
| Ras-related protein Rab-35 | Q15286 | **0.0002** |

**^1.^** Determined by ANCOVA adjusting for age and sex

**^** Variance homogeneity or approx. equal sample size were evaluated for appropriateness of ANCOVA testing. These proteins did not meet this assumption.

**Supplementary Table 6 –Proteins with increased abundance in the plasma of Recovered donors compared to Healthy donors**

| **Protein** | **UniProtKB accession** | **p-value^1^** |
| --- | --- | --- |
| Apolipoprotein C-I | P02654 | 0.0317^ |
| Immunoglobulin heavy variable 1-46 | P01743 | 0.0426 |
| Apolipoprotein B-100 | P04114 | 0.0025^ |

**^1.^** Determined by ANCOVA adjusting for age and sex

**^** Variance homogeneity or approx. equal sample size were evaluated for appropriateness of ANCOVA testing. These proteins did not meet this assumption.

**Supplementary Table 7 –Proteins with decreased abundance in the plasma of Recovered donors compared to Healthy donors**

| **Protein** | **UniProtKB accession** | **p-value^1^** |
| --- | --- | --- |
| Profilin-1 | P07737 | 0^ |
| Immunoglobulin heavy constant delta | P01880 | 0.0239^ |
| Pyruvate kinase PKM | P14618 | 0.0026^ |
| Glyceraldehyde-3-phosphate dehydrogenase | P04406 | 0.0053 |
| Cofilin-1 | P23528 | 0^ |
| Beta-actin-like protein 2 | Q562R1 | 0.0015^ |
| L-lactate dehydrogenase B chain | P07195 | 0.0001^ |
| Platelet factor 4 (PF-4) | P02776 | 0.0128^ |
| Transgelin-2 | P37802 | 0^ |
| 14-3-3 protein zeta/delta | P63104 | 0^ |
| Platelet basic protein (PBP) (C-X-C motif chemokine 7) | P02775 | 0.0018^ |
| Ras-related protein Rap-1b | P61224 | 0^ |
| Talin-1 | Q9Y490 | 0.0258^ |
| Vinculin (Metavinculin) | P18206 | 0^ |
| Integrin-linked protein kinase | Q13418 | 0.0231 |
| Fermitin family homolog 3 | Q86UX7 | 0.0001^ |
| Alpha-actinin-1 | P12814 | 0.0004^ |
| Ras-related protein Rab-15 | P59190 | 0.0004 |
| SH3 domain-binding glutamic acid-rich-like protein 3 | Q9H299 | 0.0007 |
| Peptidyl-prolyl cis-trans isomerase A | P62937 | 0.0001^ |
| Triosephosphate isomerase (TIM) | P60174 | 0.0079 |
| Ras-related protein Rab-35 | Q15286 | 0.0007 |

**^1.^** Determined by ANCOVA adjusting for age and sex

**^** Variance homogeneity or approx. equal sample size were evaluated for appropriateness of ANCOVA testing. These proteins did not meet this assumption.
